# *In vitro*-prepared A30P alpha-synuclein fibrils adopt the conserved and disease-relevant Greek key fold

**DOI:** 10.64898/2026.02.03.703603

**Authors:** Moses H. Milchberg, Owen A. Warmuth, Collin G. Borcik, Ruixian Han, Benjamin D. Harding, Dhruva D. Dhavale, Paul T. Kotzbauer, Elizabeth R. Wright, Charles D. Schwieters, Chad M. Rienstra

## Abstract

The pathological hallmark of Parkinson Disease (PD) is the formation of the protein alpha-synuclein (Asyn) into β-sheet rich, self-templating fibrils in the brain. Since the first atomic structure of wild-type Asyn fibrils was determined nearly a decade ago, several other *in vitro* structures of hereditary mutant fibrils and structures derived from post-mortem diseased patient tissue have been determined by solid-state nuclear magnetic resonance (SSNMR) spectroscopy and cryo-electron microscopy. These structures have not only expanded the library of structures available for computational modeling of drug binding and therapeutics development but have also given unprecedented insight into the disease specificity and structural polymorphism of Asyn fibrils. Here, we report the high-resolution SSNMR structure of the A30P hereditary mutant Asyn fibril, associated with early-onset PD. Our structural model is calculated using several thousand distance restraints derived from one sample, primarily sourced through 3D ^13^C-^13^C-^13^C correlation experiments. The structure adopts a Greek key topology yet does not include the P30 mutation site within the fibril core. We also introduce a comprehensive method for the rapid comparison of SSNMR spectra between Asyn polymorphs of known structure and validate the A30P fold. Lastly, we find that the structure is highly similar to many other experimental structures of both *in vitro* and *ex vivo* Asyn fibrils, including those with other hereditary point mutations, suggesting a conserved accessible fold.

## INTRODUCTION

Alpha-Synuclein (Asyn) is a 140-residue protein found throughout the central nervous system and largely present in the presynaptic termini of dopaminergic neurons in the brain. While its normal physiological function has remained elusive, it is involved in the regulation of the pool of synaptic vesicles at the presynaptic termini of neurons^6^. In disease, it can misfold, accumulate and form fibrillar aggregates in the brain. Such diseases are called Synucleinopathies, and consist of Parkinson Disease (PD), Lewy Body Dementia (LBD), Multiple System Atrophy (MSA) and other hereditary and sporadic disorders^7^. The pathological hallmark of PD is the deposition of Asyn aggregates in Lewy bodies (LBs) and Lewy Neurites (LNs). Clinically, PD is a progressive neurological disorder that manifests as a combination of muscle tremors, bradykinesia and other motor symptoms as a result of dopaminergic neuron death, eventually cascading into widespread neurodegeneration and death^8^. The role of Asyn in PD has been confirmed by the existence of duplications^9^, triplications^10^ and several familial point mutations – such as A18T, A29S^11^, A30P^12^, A30G^13^, E46K^14^, H50Q^15^, G51D^16^, A53T^17^, A53E^18^, E83Q^19^ –in the SNCA gene which encodes the Asyn protein, many of which are associated with earlier onset and increased severity of clinical and pathological symptoms. While the vast majority of PD cases are idiopathic, PD caused by a hereditary Asyn mutant presents a window into the distinct pathways Asyn might take to aggregate into fibrils and cause disease.

A30P is a hereditary mutation in Asyn associated with classical PD, first discovered in a German patient^12^ who displayed symptoms similar to idiopathic PD. Autopsy revealed that this brain tissue exhibited a much higher load of insoluble fibrillar aggregates than typical idiopathic subjects, strongly implicating fibrillar Asyn in the PD pathogenesis^20^. One hypothesis for the physiological role of Asyn involves its role in membrane plasticity, promotion of SNARE complex assembly, and binding and clustering synaptic vesicles to promote Ca^2+^-triggered exocytosis^6, 21, 22^. Solution Nuclear Magnetic Resonance (NMR) studies have shown that Asyn binds micelles and lipid vesicles through several N-terminal KTKEGV residue repeats^23-26^. Upon binding, wild type (WT) Asyn residues V3-V37 and K45-T92 form highly helical conformations^27^. Studies have shown that the A30P mutation disrupts the first helix, decreasing the binding affinity for lipid vesicles relative to WT^28-31^. The mutation has also been shown to diminish the ability for Asyn to mediate synaptic vesicle clustering^32^, leading to an excess of lipid-free A30P Asyn. The higher concentration of free Asyn, along with an increased propensity to form oligomers, are proposed to increase the inclination of A30P to aggregate and form fibrils^28, 33, 34^.

Asyn fibrils are biomarkers for disease, and determining their atomic-resolution structures is a crucial step in the development of diagnostic and therapeutic tools for synucleinopathies and can give insight into fibril formation mechanisms. Over the past decade, advances in both solid-state NMR (SSNMR) spectroscopy and cryo-electron microscopy (cryo-EM) have made it possible to investigate the atomic-level structure of Asyn fibrils, both *in vitro* and of fibrils extracted from the post-mortem brains of synucleinopathy patients^35-39^. These fibrils consist of cross-beta sheet monomers templated and stacked on top of one another into long filaments. The rigid beta-sheet core spans approximately from residues 40-100 with the N- and C-terminal tails remaining dynamically disordered and referred to as the fuzzy coat. Structural polymorphism in the number and arrangement of beta-sheets within the ordered fibril core and protofilament count and orientation at the level of quaternary structure have been widely observed and are highly sensitive to buffer, pH, salt concentration, mutation and other mechanical factors^40-44^.

Prior studies using Circular Dichroism and Atomic Force Microscopy have shown that A30P Asyn fibrils have similar secondary structure and morphological features as WT Asyn fibrils^45^. Through SSNMR studies, we previously showed similarity in chemical shifts between A30P and WT Asyn fibrils, indicating similarities in secondary structure within the fibril core; however, the high-resolution atomic-level structure of A30P Asyn fibrils had until now remained undetermined^46^.

Since the time of this past study, our lab has used SSNMR to determine the atomic-level structure of both *in vitro* WT (PDB: 2N0A^35^) and *ex vivo* amplified LBD (PDB: 8FPT^47^) Asyn fibrils. These successful efforts both relied on several deuterated and skip-labeled protein samples, as well as complex heteronuclear correlation experiments (such as ^15^N-dephased, ^13^C-detected FS-REDOR) to detect long-range distances between nuclei. The convergence of a given fibril structure calculation greatly depends on the number, quality and breadth of long-range interresidue distance restraints^48^, so determining atomic-level structures for these two fibrils were monumental tasks requiring multiple samples, complex experiments and countless hours of manual analysis. Recently, significant technological developments have been made which drastically increase spectrometer sensitivity and data resolution while decreasing overall experiment time, resulting in an unparalleled access into the atomic-level structure of Asyn fibrils^49, 50^. In addition, major advances in data analysis automation have brought about a faster and more streamlined protocol for determination of atomic-level fibril structure by SSNMR^51, 52^.

Here, we use SSNMR spectroscopy to determine the atomic-level structure of *in vitro*-prepared A30P Asyn fibrils formed in phosphate buffer. We leverage recent advances in SSNMR pulse sequence development, data processing and analysis pipelines to generate several thousand distance restraints using only one uniformly-^13^C,^15^N labeled sample and 2D and 3D homonuclear recoupling spectra. By comparing the WT, A30P and other mutant structures, we identify conserved motifs and key stabilizing contacts that persist regardless of mutation and offer insight into the potential disease-relevance and fibril formation mechanism of the A30P fold.

## MATERIALS AND METHODS

### Protein Expression and Growth

Natural abundance and ^13^C, ^15^N-labeled A30P full-length Asyn monomer were expressed and purified using previously described methods^46, 53^. Briefly, recombinant A30P Asyn protein was expressed in Escherichia coli BL21(DE_3_) cells, grown in minimal medium supplemented with ^13^C, ^15^N BioExpress (Cambridge Isotopes). After incubation at 37° C to an OD600 = 0.7, expression was induced with 0.5 mM dioxin-free isopropyl-β-thiogalactopyranoside (IPTG). Protein was harvested and subsequently purified via thermal lysis, hydrophobic interaction chromatography and size exclusion chromatography.

### Fibril Formation

1 mM natural abundance A30P monomer was solubilized into fibril formation buffer (50 mM sodium phosphate, pH 7.5, 0.1 mM EDTA, 0.02% NaN_3_), filtered with a 0.22-μm syringe filter, and incubated at 37° C for three weeks while being continuously shaken at 200 rpm. This produced mature fibrils, which were then seeded with ^13^C, ^15^N-labeled A30P monomer under the same conditions (1:100 seed-to-monomer ratio) to produce uniformly ^13^C, ^15^N-labeled A30P Asyn fibrils. After the six weeks of fibril formation, the fibril solution was centrifuged to remove the supernatant, washed with buffer, dried and packed into a 3.2 mm thin wall rotor (Varian, Fort Collins, CO), and rehydrated with 36% (m/v) water according to previously described protocols^54^.

### SSNMR Data Collection

Magic angle spinning (MAS) SSNMR experiments were collected on a 17.6 T (750 MHz ^1^H frequency) Agilent Technologies VNMRS spectrometer with a 3.2-mm Balun probe (Varian). Spinning was controlled with a Varian MAS controller to 12,500 ± 5 Hz. Pulse widths were ∼2.2-2.3 µs for ^1^H, ∼2.4 µs for ^13^C and ∼4.8 µs for ^15^N. We used 80 kHz small phase incremental alteration (SPINAL) decoupling during evolution^55, 56^. Most 2D ^13^C-^13^C experiments were measured using ^1^H-^13^C cross polarization (CP)^57^ followed by a period of dipolar-assisted rotational resonance (DARR) recoupling^58^. 2D ^15^N-^13^Cα and ^15^N-(^13^Cα)-^13^CX spectra were measured using both ^1^H-^15^N CP followed by ^15^N-^13^C’ or ^15^N-^13^Cα SPECIFIC CP^59^ and 50 ms of ^13^C-^13^C DARR mixing for the latter experiment.

3D dipolar correlation spectra were used to measure the backbone and sidechain resonances of rigid residues. Specifically, we measured 3D ^13^Cα-^15^N-^13^C’, ^15^N-^13^Cα-^13^CX, ^15^N-^13^C’-^13^CX and ^13^Cα-^15^N-(^13^C’)-^13^Cα spectra using 6 µs ^15^N-^13^C (or ^13^C-^15^N) CP contact time followed by 50 ms of DARR mixing^59, 60^. 3D ^13^C-^13^C-^13^C correlation spectra were collected with two steps of DARR mixing, one with 50 ms and another with 500 ms for the detection of long-range distance restraints. The 3D ^13^C-^13^C-^13^C correlation spectra were acquired with the 3.2-mm Balun probe in HC mode to increase the signal-to-noise ratio (SNR) by a factor of almost three compared to HCN mode (see Figure 4,^49^). These experiments were acquired with non-uniform sampling (NUS) to decrease overall experiment time^61^. NUS schedules were created using the online schedule generator from Harvard Medical School, constructed using 25% sparsity, 384 points in both indirect dimensions a front-weighted exponentially damped sine function (http://gwagner.med.harvard.edu/intranet/hmsIST/gensched_new.html).

Most experiments were performed with a variable temperature setting of 0 °C. Some 2D ^13^C-^13^C and ^15^N-^13^Cα experiments were collected with a variable temperature setting of –50 °C as a means to freeze out the molecular motions of non-rigid residues and obtain residue-type assignments. Chemical shifts were externally referenced using the downfield peak of adamantane at 40.48 ppm^62^. Chemical shifts for this sample were confirmed to largely agree with those previously reported in for A30P Asyn fibrils formed in phosphate buffer^46^, with 2D ^13^C-^13^C, 3D ^15^N-^13^Cα-^13^CX and 3D ^13^Cα-^15^N-(^13^C’)-^13^CX correlation experiments.

Spectra were processed in NMRPipe with back linear prediction, apodized using a phase-shifted cosine bell, and zero filled in the time domain before Fourier transformation and phasing^63^. Spectra were extended in the indirectly detected dimensions using Sparse Multidimensional Iterative Lineshape Enhanced (SMILE)^64^ before being analyzed in NMRFAM-Sparky at 6x the noise floor with a 1.25 x multiplication factor between contour levels^65^. Sequential resonance assignment was completed using established protocols^66^.

### NMR Datasets and Peak Picking

Datasets used for automated distance restraint generation and subsequent structure calculation were the 2D ^13^C-^13^C and 3D ^13^C-^13^C-^13^C correlation spectra (**Table S1**). Peaks in the 2D ^13^C-^13^C spectra were picked manually within NMRFAM-Sparky. Peaks in the 3D spectra were picked using restricted peak picking to avoid any spectral or SMILE reconstruction artifacts, using those from the ^13^C-^13^C 50 ms DARR spectrum with a 0.3 ppm tolerance in each dimension.

### Xplor-NIH Structure Calculation

Peaks from multidimensional spectra were subjected to automated distance restraint assignment using the Probabilistic Assignment Algorithm for Automated Structure Distance cutoffs were binned based on peak intensities as strong, medium, weak and very weak, corresponding to upper limits of 3.0, 4.0, 5.0 and 6.0 Å for the 50 ms DARR data; 4.0, 5.0, 6.0 and 8.0 Å for the 125 ms DARR data; and Determination (PASD) for fibrils^52^. Briefly, peak assignments, shift assignments and repulsive distance restraints, were generated for each of the five peak lists (**Table S1**; experiments 5-7, 15-16) under the simplifications imposed by strict symmetry. Tight chemical shift tolerances of 0.25 ppm were used for each ^13^C dimension, matching peaks to resonance frequencies in the resonance list (**Table S2**; BMRB accession code 31269). 4.0, 6.0, 8.0 and 10.0 Å for the 500 ms DARR data.

After the initial matching, the network filter was run as described previously^52^, with the following important exception. All peaks with at least one *intra*residue, sequential or short-range (|i-j|≤ 2) peak assignment were not removed at this stage and kept as distance restraints for the subsequent passes.

The PASS 2 and PASS 3 structure calculations were performed as described previously, with the starting protomer truncated to residues 20-113 to include the P30 mutation site and residues into the N- and C-termini. In addition to the automated restraints described above, TALOS-N-predicted φ and ψ backbone torsion angles (**Table S3**) and χ1 torsion angles (**Table S4**) were converted to Xplor-NIH restraints, and manual restraints were picked from unambiguous peaks in 2D and 3D datasets (**Table S5**) and inputted as separate energy terms. A vecPairOrientPot energy term was employed to restrain the fibril core as a single, non-overlapping protomer, essentially enabling the fibril to fold as a ribbon. An interprotomer NOEPot energy term was applied as well, ensuring the fibril contained the characteristic 4.7 Å spacing between protomers within a single protofilament, as is the case in cross-β sheet amyloid fibrils^67^. Calculation of 500 structures for each PASS and structural summarization was performed using the protocol detailed in **Table S6**. All energy terms used within PASD and Xplor-NIH scripts are detailed in **Table S7**. UCSF-ChimeraX was used to generate figures^68^.

### Fibril Spectrum Comparison and ZNCC Score Clustering

NMRFAM-BPHON, available as a plugin within UCSF-ChimeraX, was used to simulate spectra found in **Figure 4** and **Figure S6**^68, 69^. Simulated spectra and experimental spectra were processed with identical back-linear prediction and apodization parameters, with a phase-shifted cosine bell offset of 0.25, in NMRPipe^63^. Spectra were plotted in NMRFAM-Sparky^65^.

For experimental spectra compared in **Figure 5**, spectra were processed with identical parameters in terms of back-linear prediction, apodized and zero-filled to twice the size of the indirect dimension and extracted such that the spectra were the same size in cases where the spectral widths were different between different samples. Spectra were processed within NMRPipe^63^ and converted to text files using an in-house tcl script. The back-end of NMRFAM-BPHON^69^ as well as in-house command-line scripts were used to calculate the pairwise zero-normalized cross correlation (ZNCC) scores between each spectrum’s text file equivalent. The pairwise score matrix was then used as input into an in-house R script to perform hierarchical clustering on the ZNCC scores and plot them in a heatmap. Dendrograms are plotted above the heatmap to cleanly display which spectra cluster with one another based on similarity of pairwise scores.

### Structural Alignment

Fibril structure PDB files (PDBs) were gathered in UCSF ChimeraX^68^ via searching the RCSB Protein Databank^70^. PDBs were aligned, clustered, overlaid and analyzed using the procedure outlined in Milchberg et al.^4^. Briefly, structures were aligned using MUltiple Sequence Comparison by Log-Expectation (MUSCLE)^71^ to make their Cα coordinates invariant to one another. Most of the structures contained continuous fibril cores spanning residues V40-K96, so their coordinate files were truncated to only include those coordinates in the subsequent analysis. Principal component analysis (PCA) was performed on the aligned PDB files and DBSCAN clustering was employed on the PCA coordinates, all using functions found within the Bio3D library in R^72^.

## RESULTS

### Assignments and secondary structure of A30P Asyn fibrils

To make Asyn fibrils suitable for SSNMR data collection, we began by expressing and purifying uniformly ^13^C, ^15^N-labelled A30P Asyn and subjected it to shaking and incubation using previously established protocols^53^. We collected multidimensional homonuclear (^13^C-^13^C) and heteronuclear (^15^N-^13^C) cross-polarization (CP) SSNMR spectra (**Table S1**) – which are specific for rigid residues found in the fibril core – on the fibrils to perform chemical shift assignments. Previously, we reported the chemical shift assignments on uniformly ^13^C, ^15^N-labelled A30P Asyn fibrils^46^, but improved spectrometer sensitivity and data processing workflows have enabled us to extend the assignments further into the N-terminus, up to T33, and in regions previously thought to be dynamically disordered (V55-E57, E61-V63). We collected a 2D ^15^N-^13^Cα spectrum (**Figure 1A**) which serves as a structural fingerprint, with each peak representing one residue found in the fibril core. Expansions of the threonine Cβ-Cγ2 and Cβ-Cα regions in the 2D ^13^C-^13^C with 125 ms DARR mixing spectrum, which detail correlations primarily between intraresidue and sequential residue ^13^C atoms, show a highly ordered β-sheet structure with a beta turn at V37, noted by its unusually downfield Cα shift (**Figure 1B**). Collectively, these spectra have one set of peaks, indicative of one conformation of fibrils found within the sample.

**Figure 1:**
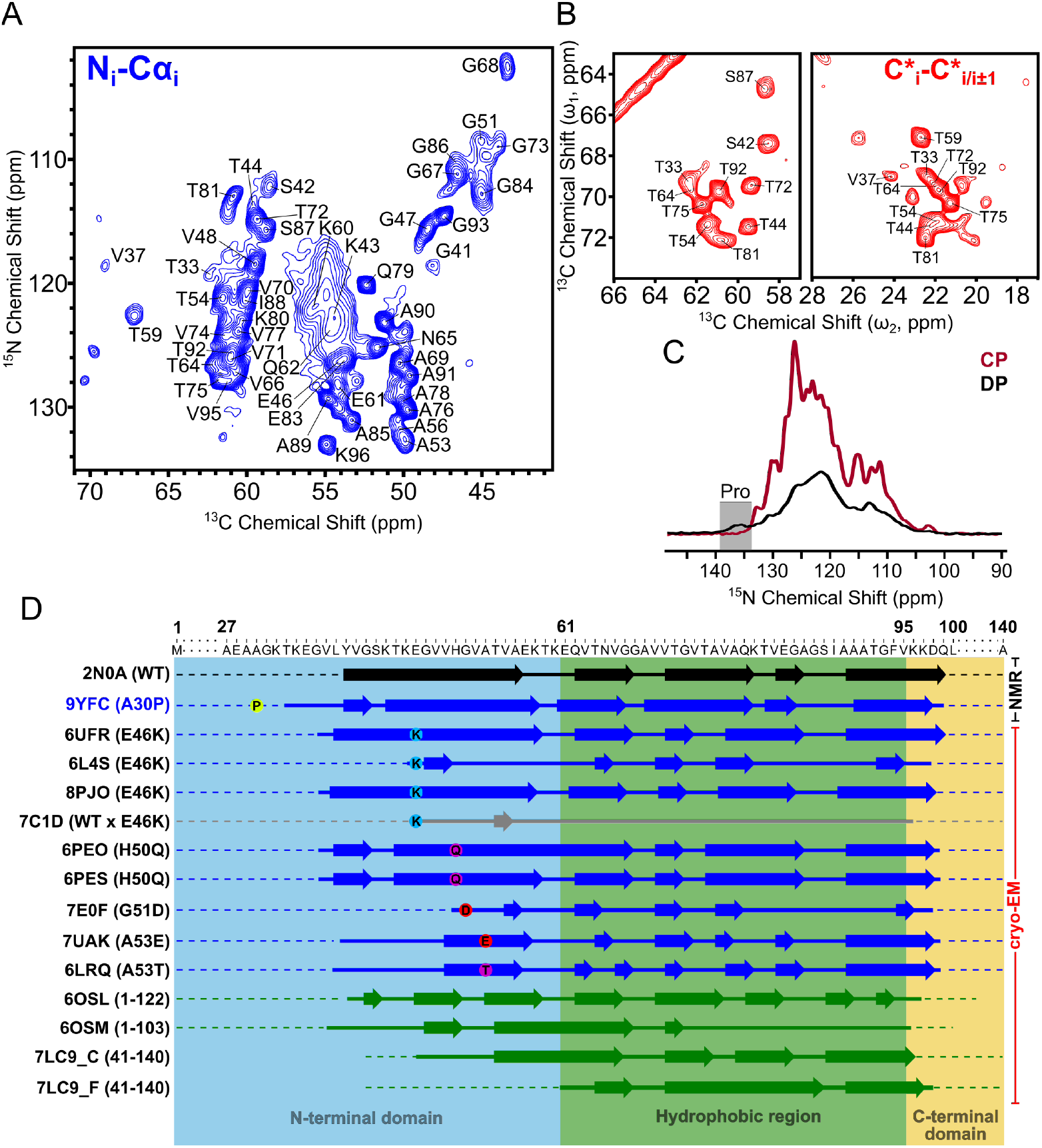
Chemical shift assignments of A30P fibrils and secondary structure comparison. (A) 2D ^15^N-^13^Ca spectrum showing correlations between backbone N[i] and Cα[i] atoms, with peaks labeled in black. (B) Portion of 2D ^13^C-^13^C with 125 ms DARR mixing spectrum showing primarily intraresidue correlations between Thr CB-CG2 in the right panel and Thr/Ser CB-CA in the left panel. (C) 1D ^15^N cross polarization (CP; brown) and direct polarization (DP; black) spectra. Peak corresponding to characteristic proline backbone ^15^N chemical shift is only present in DP spectrum. (D) Secondary structure alignment of all hereditary mutant (blue) and truncated (green) Asyn fibrils. First experimentally determined WT structure (2N0A) in black. Key regions of fibril are shaded in and labeled as such. Arrows denote regions of beta sheet structure, and thick lines indicate non-beta sheet regions that are part of the rigid fibril core. Dotted lines indicate regions of heterogeneity and/or fuzzy coat, where structure is not modeled in the cryo-EM maps. Sheet plots were made using Biotite^5^.

In order to site-specifically assign the resonances in the fibril core, we collected 3D ^15^N-^13^Cα-^13^CX, 3D ^13^Cα-^15^N-[^13^C’]- ^13^CX and 3D ^15^N-^13^C’-^13^CX spectra and assigned the shifts for each residue using standard backbone walk chemical shift assignment strategies (**Figure S2**)^58-60, 66^. We were able to site-specifically assign the backbone ^15^N, ^13^Cα and ^13^C’ resonances for each residue within the fibril core except for E35, L38, H50C’, A56C’ and D98C’, as well as 93.5% of the total carbon sidechain resonances (**Figure S1C and Table S2**). Our new assignments (BMRB deposition 31269) largely agree with the previously published assignments, with an average chemical shift difference (CSD) of 0.29 ppm (**Figure S3**). Differences above 0.3 ppm are only observed for G41, K43, V52, V55, K60 and V82, due to improved sensitivity resulting in more accurate resonance assignment of backbone ^15^N and sidechain ^13^C shifts. Using the backbone chemical shifts, we were able to predict ψ and φ torsion angles using the TALOS-N program^73^ (**Figure S1A**). We were also able to predict χ_1_ torsion angles using the Cβ and Cγ sidechain resonances for certain amino acids. The secondary structure of the fibril core consists of six beta sheets starting at Y39-G41 (β1), K43-K58 (β2), K60-V66 (β3), A69-Q79 (β4), T81-G84 (β5) and A90-K97 (β6) (**Figure S1A**). Residue stretches in between the beta strands correspond to loops and regions of local heterogeneity where sidechain resonances were difficult to site-specifically assign in the NMR spectra. The mean per-residue RCI S^2^ order parameter^74^ within the fibril core is 0.83 but decreases in areas between beta strands, indicating those regions contain increased flexibility (**Figure S1B**). 1D ^13^C spectra indicate that the fibril termini are heterogeneously disordered, specifically noted by the increase of peak intensity around 183 ppm (characteristic glutamic acid Cδ chemical shift^75^) in the direct polarization spectra; 72% of the glutamic acids within Asyn are present in the termini (**Figure S1D**).

### Fibril Core Excludes A30P Mutation Site

In our 2D ^15^N-^13^Cα CP spectrum (**Figure 1A**), we were unable to detect a peak corresponding to P30N-Cα. The ^1^H-^15^N contact time was set to 1.8 ms, which is sufficient for transferring polarization from ^1^H to ^15^N in amino acids with amide protons, but since proline does not have an amide proton, we hypothesized that increasing the contact time would allow for sufficient polarization transfer from proline ^1^Hδ2/3 or ^1^Hα to the backbone ^15^N, enabling detection of proline N-Cα or N-Cδ correlations. A30P Asyn has six prolines, one at residue 30 and the other five in the C-terminus, starting with P108, meaning that any proline resonance we see is likely P30 due to it being three residues away from T33 which is part of the fibril core. The characteristic chemical shift of the proline backbone ^15^N is roughly at 136 ppm^75^, which is significantly downfield from any other backbone ^15^N shift found in Asyn, so a resolved peak in this region would likely correspond to P30. To investigate this, we collected a series of 1D ^15^N CP spectra with varying ^1^H-^15^N contact times ranging from 1 ms to 5 ms (**Figure S4A**). While the overall signal intensity decreased with increasing contact time due to decreased polarization transfer efficiency, the intensity around 136 ppm remained unchanged (around zero). We decided to then collect a 1D ^15^N direct polarization (DP) spectrum, which directly excites all ^15^N nuclei and does not rely on dipolar couplings for polarization transfer, allowing us to detect all resonances in the fibril core *and* in the dynamically disordered termini. We were able to detect a broad peak spanning 134-139 ppm, corresponding to the P30 and C-terminal backbone ^15^N proline resonances (**Figure 1C and S4A**). In addition, we collected a 2D ^15^N-^13^Cα CP spectrum at a variable temperature set point of -50° C to freeze out the molecular motions within the fibril (**Figure S4B**). Overall, the spectrum appeared much broader than that collected at 0° C in part due to the freezing of multiple conformations within mobile regions. Peaks at a ^15^N frequency of 132-136 ppm (Proline backbone ^15^N) and ^13^C frequencies of 60-63 ppm (Proline Cα) and 49-52 ppm (Proline Cδ) were also detected albeit at low signal intensity, indicating the freezing out of mobile prolines present only at -50° C but not at 0° C. Together, these suggest that P30 remains outside of the fibril core in A30P Asyn fibrils.

Asyn can be divided into three regions: (1) the N-terminal domain, which contains the A30P mutation and several of the KTKEGV repeats serving as lipid-binding domains (M1-K60), (2) the hydrophobic region (previously referred to as the non-amyloid component or NAC)^76^ which comprises the majority of the fibril core (E61-F94) and the (C) C-terminal domain, which is highly negatively charged (V95-A140) (**Figure 1D**). To date, nine structures of hereditary point mutation fibrils (excluding A30P) and eight fibrils with sequence truncations have been determined by both SSNMR and cryo-EM and published in peer-reviewed journals. All of these mutations occur in the N-terminal domain, within the rigid fibril core (approximately V40-L100). The sequence truncations occur in both the C- and N-termini, yet do not largely affect the location of the fibril core. The A30P mutation stretches the fibril core to T33, a few residues more than other reported mutant structures. At the level of secondary structure, the placement of beta-strands is largely conserved, regardless of mutation or truncation, suggesting that sequence may restrict the folding landscape of fibrils but is not solely the determining factor in fibril secondary structure.

### Long-range Contacts Reveal the Tertiary Structure of A30P Fibrils

We collected 2D ^13^C-^13^C and 3D ^13^C-^13^C-^13^C correlation spectra on the fibril sample to detect long-range distance correlations between atoms. In the 2D spectrum, we set the DARR mixing time to 500 ms, enabling the detection of ^13^C-^13^C distances typically up to 10 Å apart^58, 78, 79^. The specific 3D homonuclear experiment we ran had two DARR mixing steps, one of 50 ms which correlates two ^13^C nuclei within ∼6 Å, followed by a mixing step of 500 ms, where magnetization is transferred to ^13^C nuclei up to ∼10 Å away from the ω_2_ ^13^C nucleus. While these spectra contain tens of thousands of peaks collectively (**Table S1**), rendering manual assignment of each peak a laborious task, we manually assigned peaks corresponding to unambiguous long-range distances, defined as peaks with only one resonance assignment possibility within a tolerance window of 0.35 ppm, or about 1.5 times the average ^13^C linewidth. These peaks consisted of multiple contacts between residues on opposing beta strands, including E46-A78/K80, V74-V52, I88-F94/K96, and G93-V71 (**Figure 2A**). These contacts are displayed on the lowest energy structure following refinement (**Figure 2B**). Additional manually assigned correlations used in the Xplor-NIH structure calculation can be found in **Table S5**.

**Figure 2:**
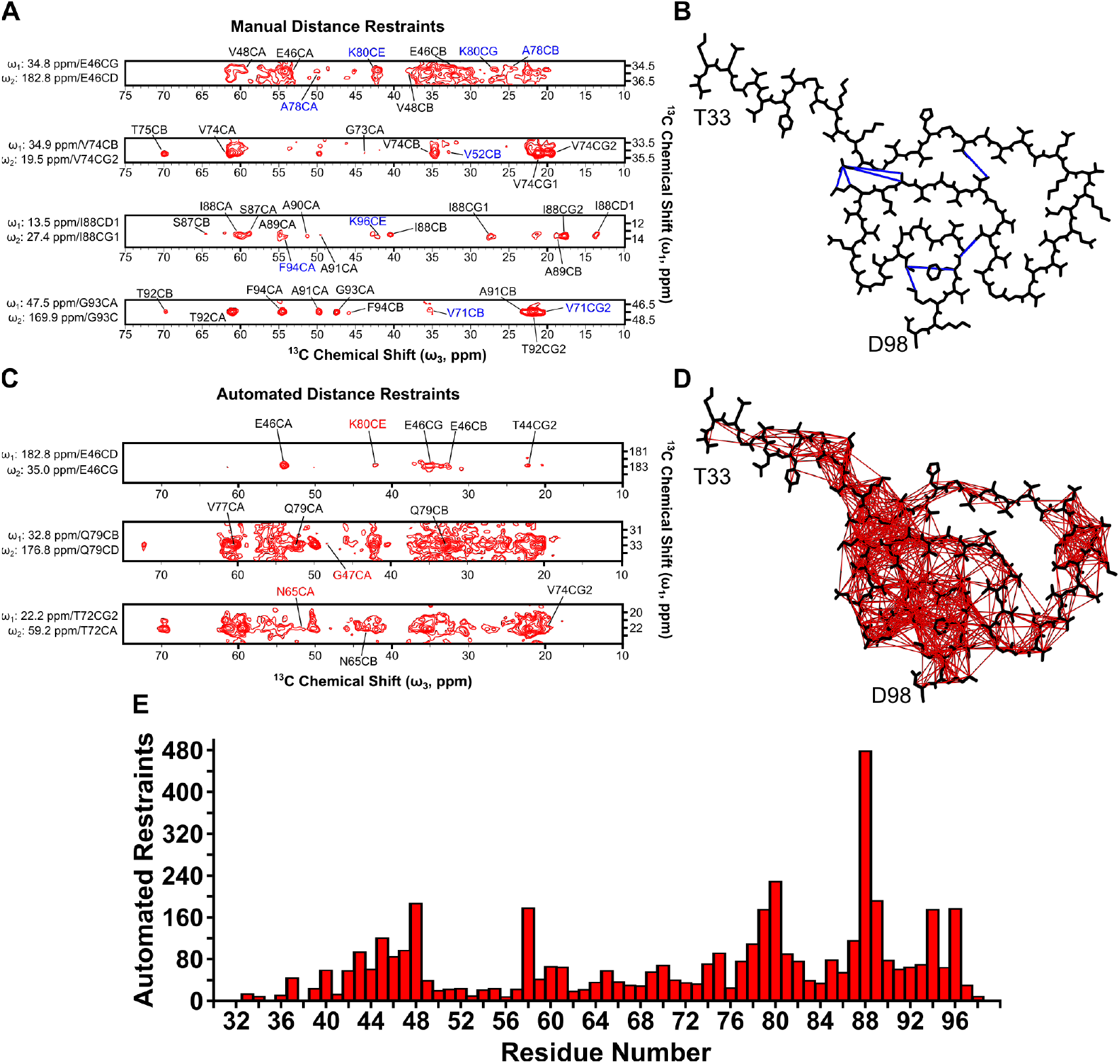
Manual and automated distance restraints derived from 3D ^13^C-^13^C-^13^C 50 ms, 500 ms DARR mixing spectrum. Representative strips with (A) unambiguous peak assignments and (C) probabilistic peak assignments derived from the automated distance restraint assignment protocol, PASD, corresponding to ^13^C-^13^C distances up to 10 Å in space. Peaks labeled in blue (A) and red (C) correspond to distance restraint assignments > 3 residues apart in the primary protein sequence. The manual distance restraints were used as a standalone energy term in the Xplor-NIH structure calculation. Lowest energy protomer after structure calculation with (B) representative manual (blue) and (D) automated (red) distance restraints overlaid. (E) Automated restraints on a per-residue basis used in the Xplor-NIH structure calculation, with an average of 67 restraints per residue.

### Structural Model of A30P Asyn Fibrils

While we were able to determine several unambiguous manual distance restraints from the 3D ^13^C-^13^C-^13^C spectrum, over 23,000 peaks were picked in the spectrum, a significant number of which are overlapped and/or ambiguous. Traditionally, a subset of these peaks would be manually picked and left as formally ambiguous in the structure calculation. Instead, we used the PASD protocol within Xplor-NIH to perform automated assignment of these peaks to attribute likelihoods to ambiguous distance restraints^52^. Because of the robustness and speed of the automated assignment in PASD, we applied the protocol to peak lists in several other 2D and 3D spectra: (1) 2D ^13^C-^13^C 50 ms DARR mixing, (2) 2D ^13^C-^13^C 125 ms DARR mixing, (3) 2D ^13^C-^13^C 500 ms DARR mixing, and (4) a second 3D ^13^C-^13^C-^13^C 50 ms, 500 ms DARR mixing spectrum (**Table S1**).

To calculate the structure of A30P Asyn fibrils, we used the PASD-defined correlations to generate distance restraints within Xplor-NIH^51, 52, 80^. The strict symmetry facility was employed to model just the protomer, using rotational and translational symmetry to model a 5-subunit single protofilament fibril. A summary of the simulated annealing protocols used in the overall structure calculation can be found in **Table S6**, and the details of the Xplor-NIH energy terms used can be found in **Table S7**. Violation statistics for high-likelihood restraints for the final structural bundle after the final refinement can be found in **Table 1**. Given the resonance list for the ^13^C-detected nuclei, the initial matching (initMatch) stage in PASD resulted in 18885 cross-peaks from data collected on homonuclear ^13^C correlation spectra, with 6984 of the cross-peaks corresponding to assignments where |i-j| ≥ 5 (between residues separated by at least 5 residues in primary sequence). 4308 ± 6, or 23% of these peaks contained assignments which were satisfied by the final ensemble of 10 structures. The initial matching resulted in a weighted average ambiguity of 35 ± 43 peak assignments per peak. The network analysis stage (jointFilter) reinforced assignments by giving potential peak assignments between a pair of residues higher weights when multiple such assignments exist. Additionally, at this stage peaks with any protomer ambiguity of greater than two were removed, and all peaks which have more than 20 possible peak assignments were deleted, resulting in a reduction in the overall ambiguity to 7 ± 7. Three rounds of PASD structure calculations (PASS 2, PASS 3 and PASS 4) removed nearly all the violating restraints while adding back in the satisfied restraints, following the protocol outlined previously^52^.

**Table 1.**
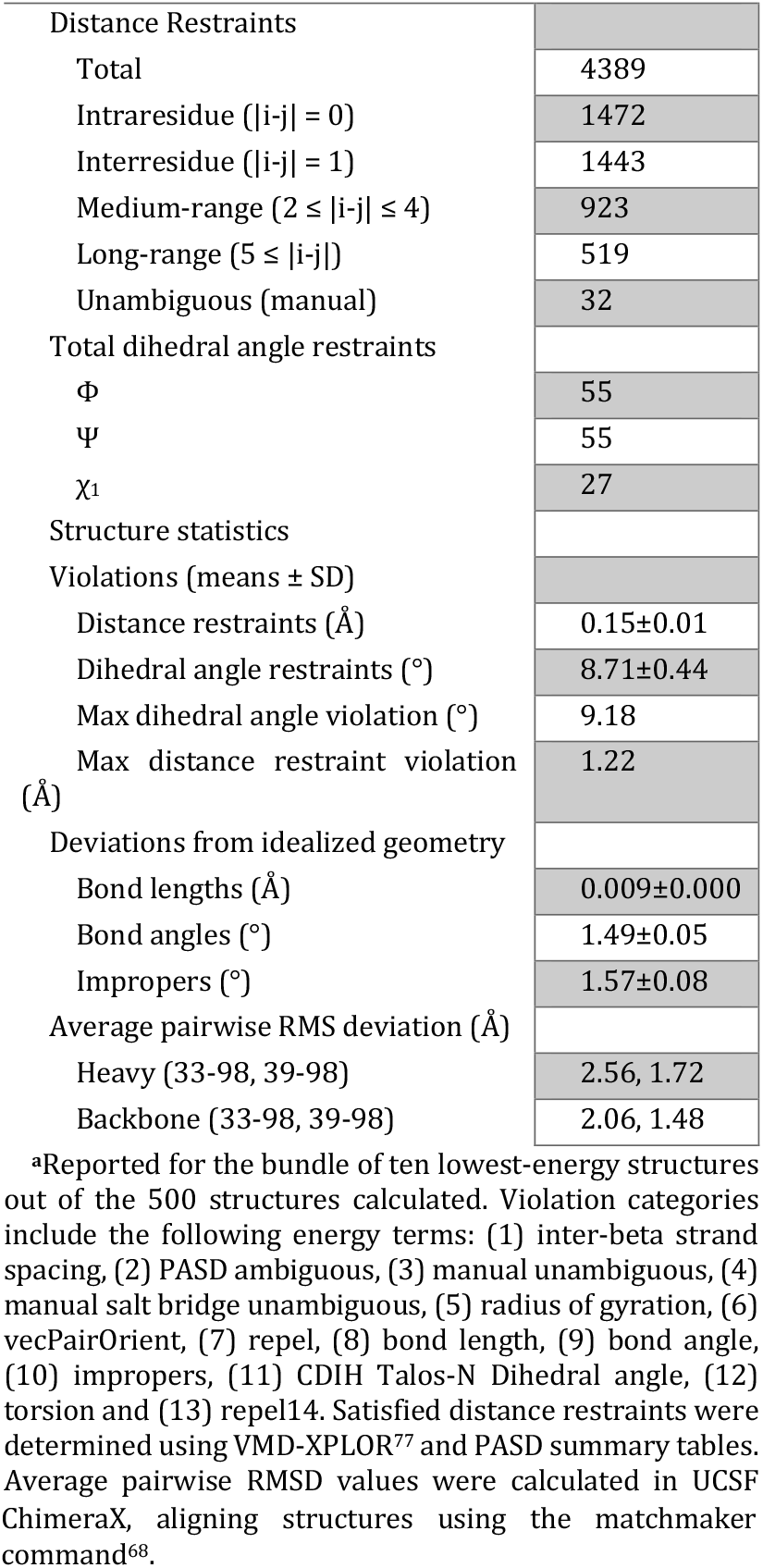
NMR Distance and Dihedral Restraints^a^.

The structural model of A30P Asyn fibrils is shown as the protomer of the lowest energy structure in **Figure 3A** and as an overlay of the ten lowest energy structures in **Figure 3B**, with residues T33-D98 modeled. The structure contains canonical features of Class 1A Asyn fibrils^4^, such as a (1) Greek key motif between residues A69-V95 with close interaction of the I88, A91 and F94 sidechains (**Figure 3C**), (2) steric zippers between G47-A78-V49-A76 and V71-G93-A69, (3) a glutamine ladder at Q79 and (4) an intermolecular salt bridge between E46-K80 (**Figure 3D**). The ensemble of ten lowest energy structures (**Figure 3B**) has a heavy atom RMSD of 1.72 Å and backbone RMSD of 1.48 Å for residues Y39-D98. For the stretch of residues between T33-L38, the lack of long-distance restraints connecting them to another beta-strand within the sequence plus lack of assignments (and therefore no dihedral angle restraints) for E35 and L38 results in a heterogeneous structural distribution. The bundle of the ten lowest energy structures has been deposited to the RCSB PDB with accession code 9YFC.

**Figure 3:**
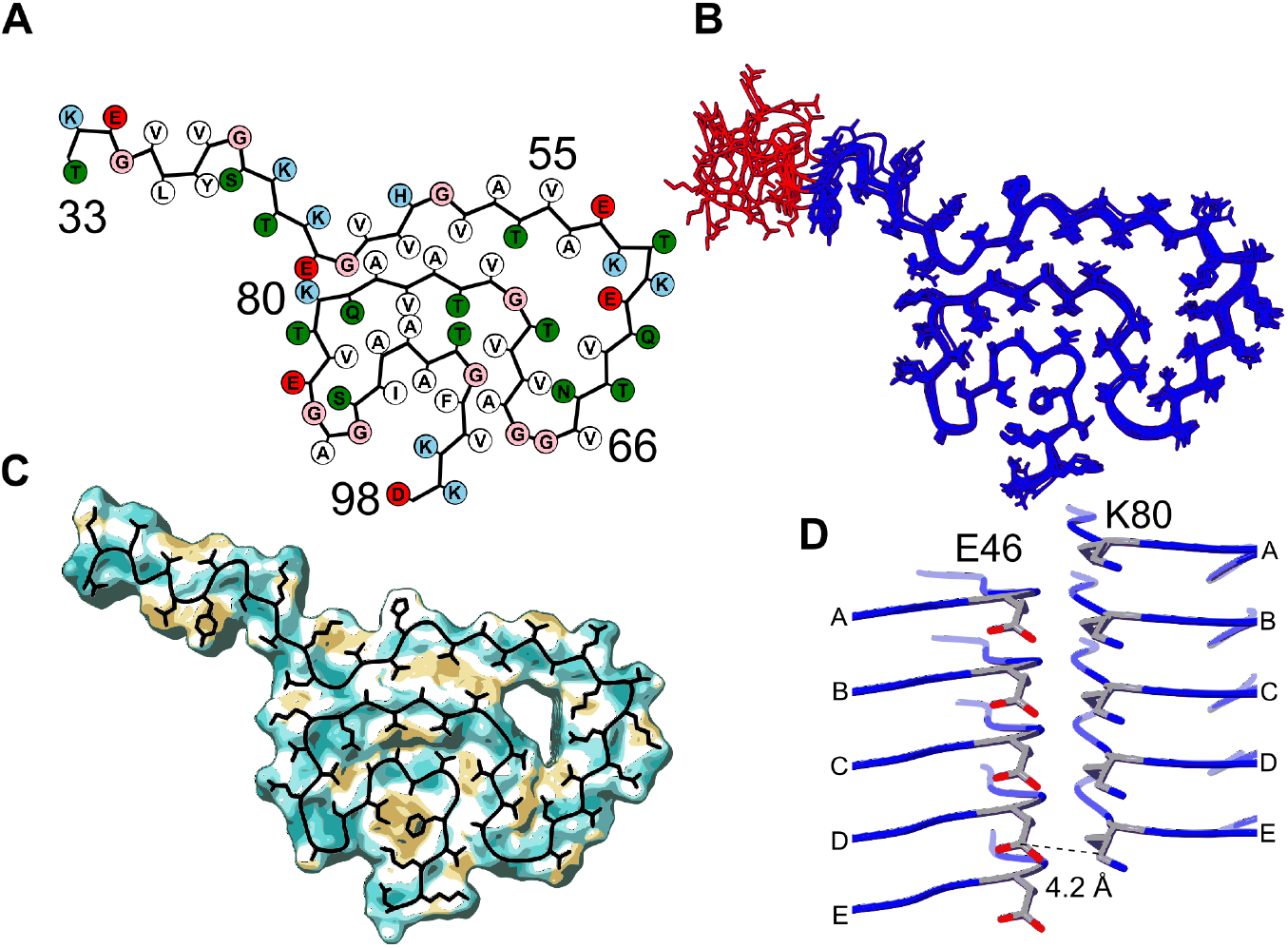
Structure of A30P Asyn fibrils. (A) Graphical representation of A30P Asyn protomer made using the atom2svg.py script^1^. Negatively charged residues are colored in red, positive in sky blue, polar in green, glycines in tan and hydrophobic in white. (B) Overlay of 10 lowest energy protomers following final refinement in Xplor-NIH structure calculation, residues Y39-D98 colored in blue with a heavy atom RMSD of 1.72 Å and backbone RMSD of 1.48 Å. (C) Molecular lipophilicity map potential^2, 3^ for A30P Asyn fibrils, with lowest energy protomer overlaid in black. The units of lipophilicity are measured in logP(octanol-water), with brighter values being more positive and hydrophobic and darker values being more negative and hydrophilic (lipophobic). The I88-A91-F94 sidechain pocket is highly hydrophobic. (D) Intermolecular salt bridge between E46Cδ and K80Cε, with a distance of 4.2 Å shown between chains D and E. The structural bundle is available on the RCSB PDB with accession code 9YFC.

Following structure determination, we can see that from the original 18885 crosspeaks detected in our spectra, 4357 automated distance restraints were generated (**Table 1**). These restraints are overlaid on the lowest energy protomer (**Figure 2D**) alongside several strips from the 3D ^13^C-^13^C-^13^C spectrum detailing a few examples of the assignment of ambiguous and/or overlapped distance restraints (**Figure 2C**). Such contacts are between E46Cγ-K80Cε, Q79Cδ-G47Cα and T72Cα-N65Cα. There is sufficient coverage of automated restraints throughout each residue within the fibril, except E35 and L38, with an average of 67 total and 8 long-range automated restraints per residue (**Table 1 and Figure 2E**). This coverage is more than enough to derive an accurate model structure (as shown by Russell and colleagues)^48^.

### Validation of Structure

In order to validate our experimental SSNMR structure of A30P Asyn fibrils, we used the program NMRFAM-BPHON, developed by Harding and colleagues^69^. This structural validation program works by taking (1) an experimentally determined structural model and (2) a resonance list and uses spin physics to back-calculate and simulate NMR spectra for direct comparison to experimental spectra. The two spectra are treated as images and compared via zero-normalized cross correlation (ZNCC) scores, where a score of 0 corresponds to complete anticorrelation and 1 corresponds to complete agreement.

We simulated 2D ^15^N-^13^Cα (**Figure S6A**), 2D ^13^C-^13^C with 50 ms DARR mixing (**Figure 4A**) and 2D ^13^C-^13^C with 500 ms DARR mixing (**Figure S6B**) spectra and calculated their ZNCC scores with respect to their equivalent experimental spectra (**Table S1**). In principle, a ZNCC score of 1 corresponds to a perfect structural model, a complete and accurate resonance list, and relative intensities and linewidths reflective of the true chemical environment of each NMR-active site in the protein. In practice, the scores – especially for fibrils – tend to be much lower than 1 due to the imperfections in the spin physics model used, the inherent resolution of the experimental structure, difficulty replicating experimental artifacts in simulated spectra, lack of incorporation of site-specific dynamics measurements, and other related reasons. Therefore, the combination of scores, relative overlap of peaks, and improvement compared to simulated spectra using a SHIFTX2-predicted resonance list can be used to assess structure validity^81^.

**Figure 4:**
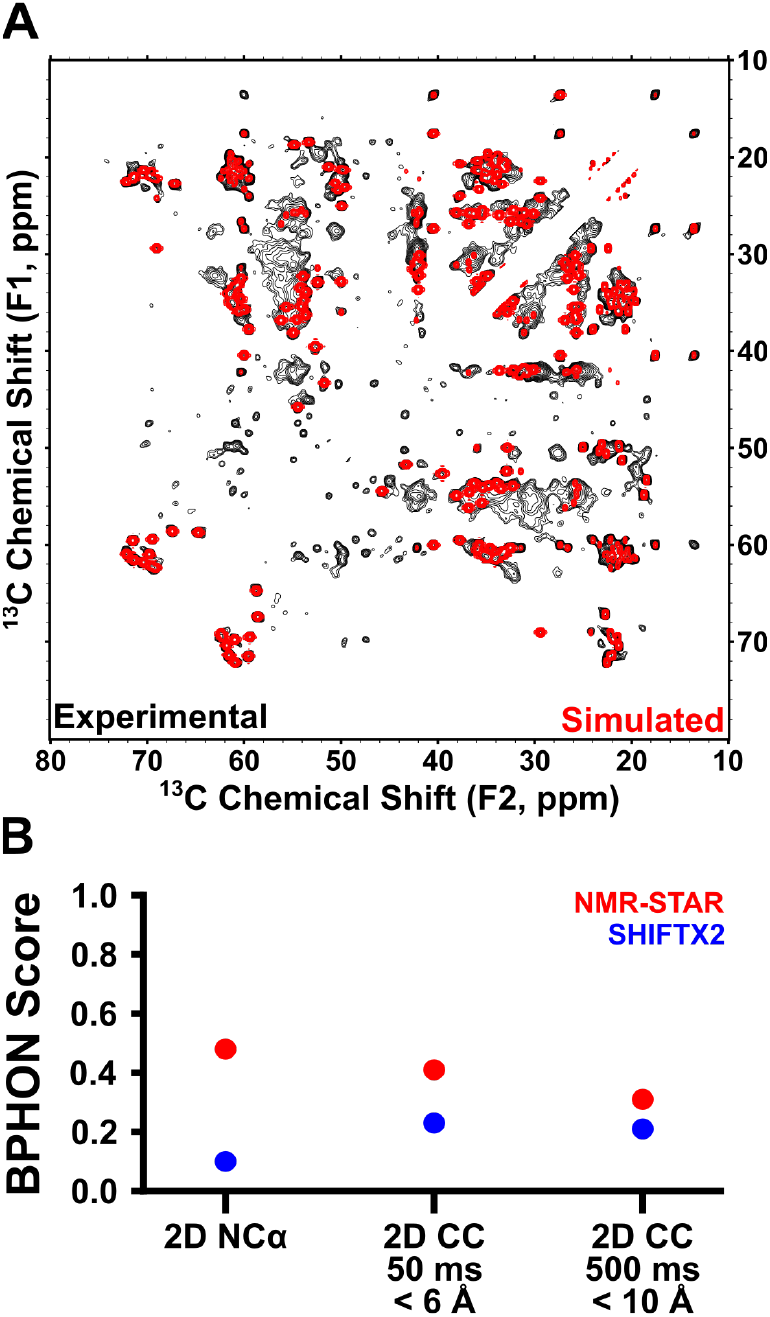
Validation of structure. (A) Overlay of experimental (black) and NMRFAM-BPHON simulated (with experimental resonance list, BMRB: 31269) (red) 2D ^13^C-^13^C with 50 ms DARR mixing spectrum, with interatomic distances up to 6 Å apart. (B) Plot of zero-normalized cross correlation (ZNCC) scores between (1) experimental spectra and NMR-STAR resonance list-derived simulated spectra (BMRB: 31269) (red) and (2) experimental spectra and SHIFTX2-predicted resonance list-derived simulated spectra (blue), for the 2D ^15^N-^13^Cα spectrum (**Figure S6A**), the 2D ^13^C-^13^C with 50 ms DARR mixing (**Figure 4A**), and the 2D ^13^C-^13^C with 500 ms DARR mixing (Simulated spectra with 200 ms mixing; **Figure S6B**), which has peaks corresponding to interatomic distances up to 10 Å apart.

We started by scoring experimental spectra versus spectra simulated using the experimental resonance list (BMRB: 31269). Here, the ZNCC scores for the 2D ^15^N-^13^Cα, 2D ^13^C-^13^C with 50 ms DARR mixing and 2D ^13^C-^13^C with 500 ms DARR mixing were found to be 0.48, 0.41 and 0.31. As a negative control, we compared experimental spectra to spectra simulated using only a SHIFTX2-predicted resonance list. In this case, the ZNCC scores for the 2D ^15^N-^13^Cα, 2D ^13^C-^13^C with 50 ms DARR mixing and 2D ^13^C-^13^C with 500 ms DARR mixing were found to be 0.10, 0.23 and 0.21, respectively (**Figure 4B**).

**Figure 4A** shows the overlay of the experimental 2D ^13^C-^13^C 50 ms DARR mixing spectra with the corresponding NMRFAM-BPHON simulated spectra. While the ZNCC score of 0.41 is much lower than 1.0, there is significant overlap throughout, particularly in the Thr Cβ-Cα and Cβ-Cγ2, Val Cβ-Cγ*, Cα-Cβ and Cα-Cγ*, and Ala Cα-Cβ regions. Peaks which are not found in the simulated spectrum include those near ω_1_ = 55 ppm and 30 ppm ≤ ω_2_ ≤ 45 ppm, which could correspond to lysine sidechain correlations. All the lysines between T33-D98, except for K34, have the majority of their sidechains assigned. In total, Asyn has 15 lysines, and seven of them appear in the heterogenous N- and C-termini, which are largely invisible to CP-type experiments. Still, some of the six N-terminal lysines and one C-terminal lysine might form salt bridges with the many N- and C-terminal glutamic acids which appear in different conformations, contributing to their rigidity and appearance in CP spectra. Although these discrepancies between the spectra exist, there are no simulated peaks which do not overlay with experimental peaks. This suggests that the shortcomings reflected in both spectral overlay and ZNCC score are primarily due to the existence of local structural order in the termini not accounted for in the chemical shift assignments.

Similarly, for the 2D ^15^N-^13^Cα spectrum, **Figure S6A** shows the overlay of the resonance-list simulated spectrum with the experimental spectrum. The ZNCC score here is 0.48, much lower than 1, but it is clear from the overlay that this discrepancy is largely due to the intensities and linewidths not being accurately reflected in the spectral simulation. The simulated peak positions all line up extremely well with the experimental data, indicating accuracy in our resonance list. There are also some regions in the experimental spectrum where there are no simulated peaks, reflecting spin systems we were either (a) unable to resolve in the 3D spectra or (b) able to residue-type assign but not fit anywhere in the linear amino acid sequence. For the 2D ^13^C-^13^C with 500 ms DARR mixing spectrum, Figure 6B shows the overlay of the resonance-list simulated spectrum with the experimental spectrum. The ZNCC score of 0.31 is also lower than one might expect, due to lower accuracy of the polarization transfer model at longer DARR mixing times. That being said, there is still significant peak overlap, consistent with the accuracy of the long-range distance restraints present in the A30P Asyn fibril structural model.

We also utilized the program SSD-NMR to predict secondary structure elements as a means of validating the extent of the A30P fibril core. SSD-NMR is able to predict the percentage of a protein’s conformation that exists as a random coil, an α-helix or a β-sheet, using the protein sequence and an experimental 1D ^13^C spectrum as inputs^82^.

We simulated both CP (**Figure S7A**) and DP (**Figure S7B**) spectra for Asyn sequences T33-D98 – corresponding to the extent of our chemical shift assignments – and for the entire sequence of M1-A140. The extent of secondary structure conformations are shown in **Figure S7C**. We can see that for the CP spectrum, assuming our fibril core spans from T33-D98, 84% of the residues are predicted to have β-sheet character. This value agrees very well with the value predicted by TALOS-N directly from our resonance list, where 84% of the residues have predicted β-sheet character (**Figure S1B**). As we extend the sequence to M1-A140, we see the β-sheet character decrease to 80% and the α-helical character increase from 11% to 15%, indicating that the M1-K32 and Q99-A140 tails are not β-sheets. For the DP spectra, loop regions – which are accounted for in 11% of the fibril core residues (**Figure S1B**) – can equally contribute along with β-sheets to the peak intensities in the spectrum, explaining the significant increase from 5% to 27% coil character between the CP and DP T33-D98 SSD simulations (**Figure S7C**). In turn, the DP β-sheet character decreases to 60%. These spectral simulations and predictions of secondary structure agree well with our experimental model and serve as a complementary validation along with NMRFAM-BPHON.

### Quantitative Comparison of Asyn Fibril NMR Spectra Enables Rapid Polymorph Classification

While a structure can be validated by comparing its simulated spectra directly to the raw experimental data, we can also compare its NMR spectra to NMR spectra of other Asyn fibril samples of known structure. Previously, we have determined structures from four other Asyn fibril samples, (1) *in vitro* wild-type Asyn fibrils prepared in phosphate buffer (PDB: 2N0A)^35^, (2) *in vitro* wild-type Asyn fibrils prepared in a high ionic strength Tris buffer (PDB: XXXX)^83^, (3) *amplified LBD* Asyn fibrils derived from postmortem human LBD brain tissue (PDB: 8FPT)^47, 84^, and (4) *in vitro* wild-type Asyn fibrils prepared in phosphate buffer for an extended incubation period (PDB: 9CK3)^85^. Each of these samples has corresponding 2D and 3D SSNMR spectra which were used to determine and/or validate a structure, so we can compare a few of their representative spectra and thus compare structure.

**Figure 5A** shows an overlay of the 2D ^15^N-^13^Cα spectra of each of the four aforementioned samples as well as for A30P Asyn fibrils. The NCα spectrum serves as a structural fingerprint where the number of peaks represents the number of residues which make up the fibril core, and the position of peaks is highly dependent on secondary structure. Therefore, overlapped peaks between two representative spectra indicate similar secondary structure. We see that there is significant, but not full, overlap between the WT Tris, 2N0A and A30P samples in the NCα spectral overlay. To better quantify their similarity, we converted each spectrum to an image, calculated their pairwise ZNCC scores and clustered them into a dendrogram (see Materials and Methods) (**Figure 5B**). Each autocorrelation has an expected ZNCC score of 1.00, and spectra that have pairwise scores of less than 0.50 are not considered similar in structure. The WT Tris, 2N0A and A30P NCα spectra have pairwise ZNCC scores all above 0.80, indicating a very high degree of similarity at the level of secondary structure, again shown by similarity between their cartoon structures in **Figure 5C**.

**Figure 5:**
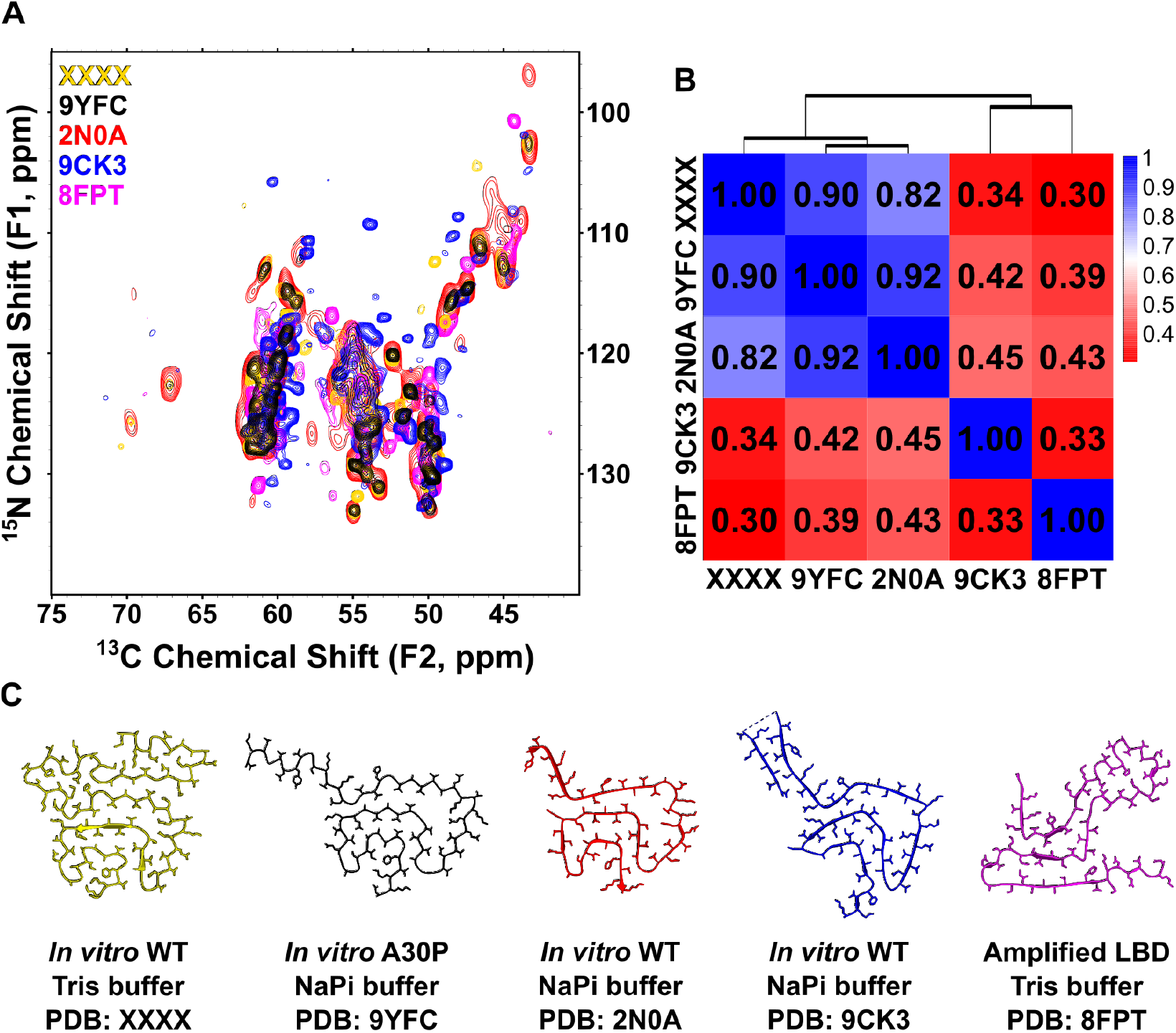
Spectral comparison of Asyn fibril polymorphs of known structure. (A) 2D ^15^N-^13^Cα spectra of *in vitro* WT fibrils formed in Tris buffer at moderate ionic strength (PDB: XXXX; yellow), *in vitro* A30P fibrils (PDB: 9YFC; black), *in vitro* WT fibrils (PDB: 2N0A; red), *in vitro* WT fibrils formed with extended incubation time (PDB: 9CK3; blue) and amplified LBD fibils (PDB: 8FPT; magenta). Similarities in NCα spectra indicate similarities in secondary structure at backbone-site resolution. (B) Pairwise ZNCC scores between each spectrum, hierarchically clustered into a dendrogram. High scores between WT Tris, 9YFC and 2N0A indicate structural similarity. (C) Protomer structures of WT Tris, 9YFC, 2N0A, 9CKS and 8FPT shown side-by-side for visual comparison.

### The A30P Fold in the Context of Asyn Fibril Structural Polymorphism

Mutations in Asyn leading to disease only account for < 1% of all synucleinopathy cases, but they offer a window into what might drive disease onset and rapid progression. The atomic-level structures of hereditary point mutation Asyn fibrils have been determined by cryo-EM, specifically for E46K^86^, H50Q^87^, G51D^88^, A53T^89^ and A53E^90^, and published in peer-reviewed journals. Additionally, several structures featuring both deletions (7LC9^91^, 6OSL and 6OSM^92^) and insertions (8BQW, 8BQV, 8CE7 and 8CEB^93^) have been determined by cryo-EM and also published in peer-reviewed journals. In a similar approach to what we employed in a previous study^4^, we aligned each unique protomer of point mutant structures including A30P, deletion/insertion structures and the first-determined *in vitro* WT structure (PDB: 2N0A). We reduced the dimensionality of the aligned structures using principal component analysis (PCA) and clustered the PC2 vs PC1 coordinates using Density Based Clustering with Applications to Noise (DBSCAN) to identify structural classes (**Figure 6A**). This revealed one major cluster of fibrils, of which A30P was included. This cluster has exact overlap to Cluster 1 from our previous study, characterized primarily by the Greek key motif between A69-V95 and an intermolecular salt bridge between E46-K80^4^. Even though 6UFR and 8PJO, both E46K mutant fibrils, formed a major cluster previously (Class 2 fibrils), they are not part of the major cluster of fibrils in this analysis. The per-residue root-mean-square fluctuation (RMSF) for each structure (between residues V40-K96) within this cluster reveals a tight similarity of < 3 Å between S42-G84 and larger differences between A85-K96, reflecting the intraclass differences dominated by the presence or absence of close interaction between the I88, A91 and F94 sidechains (**Figure 6B**). As previously shown, the increase in per-residue RMSF in this region is the key difference between Cluster 1A and 1B fibrils. Since A30P has this close interaction of hydrophobic sidechains (**Figure 3C**), it falls into Cluster 1A. The structural overlay seen in **Figure 6C** highlights the overall structural similarity but visualizes these intraclass differences, including the differences in angle at G41 which contribute to elevated localized root-mean-square fluctuation (RMSF) between V40-G41. Interestingly enough, the known small molecule binding pocket of residues K43, K45, V48 and H50^94, 95^ is well conserved across all clustered structures (**Figure 6D**). The Cluster 1A Greek key motif (close interaction of I88-A91-F94 sidechains) as well as the K43-H50 binding pocket are also found in fibrils extracted from the brains of MSA patients^38^ and *in vitro* fibrils which are capable of inducing MSA-like lesions and pathology in mouse brains (PDB: 9EUU)^96^, suggesting that the A30P fold is disease-relevant and likely induces an MSA-like pathology. Structures of Asyn fibrils of the mutations G14R^97^ and K58N^98^ have also been determined by cryo-EM, available as of now in preprints on BioRxiv, so they were excluded from this analysis.

**Figure 6:**
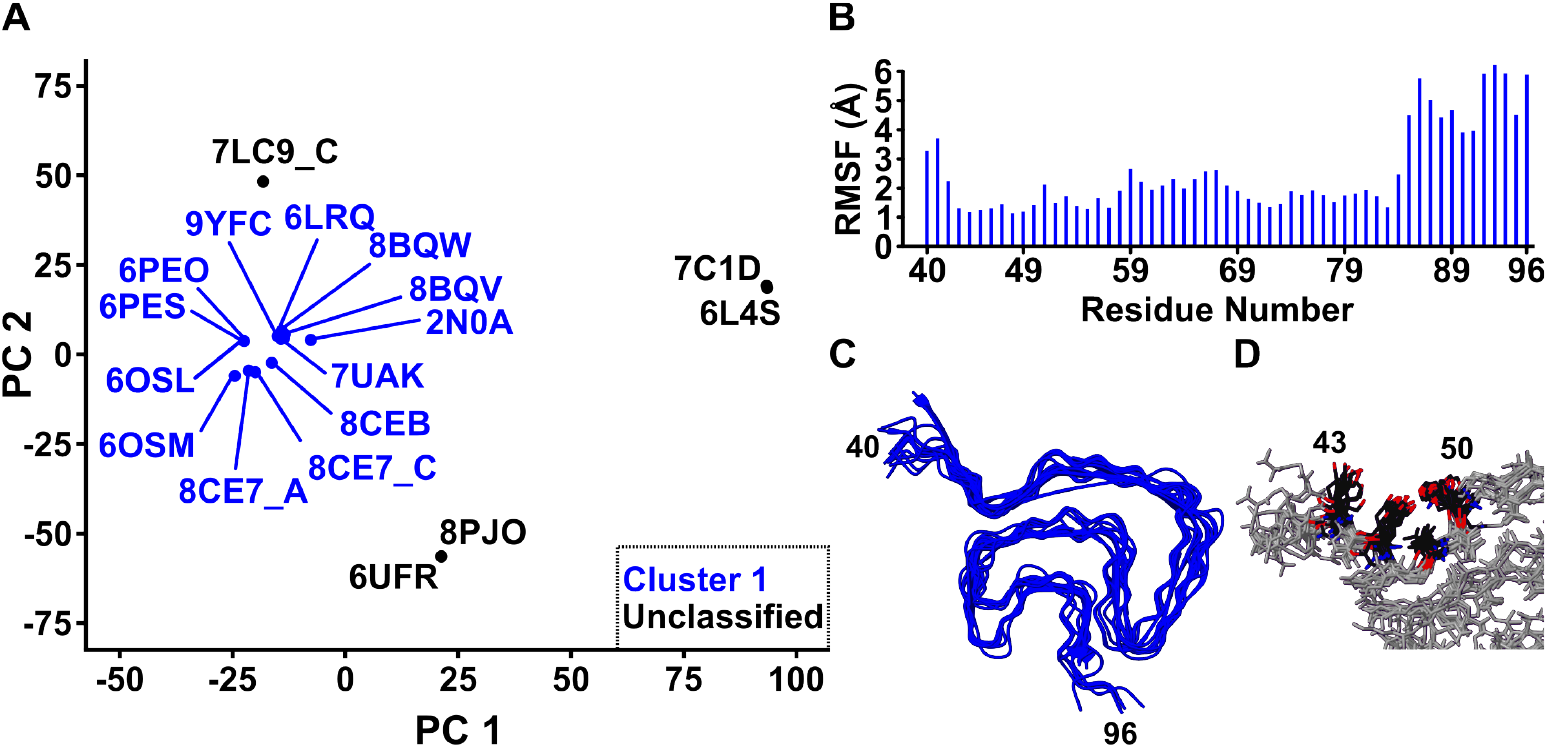
Structure comparison across Asyn fibril polymorphs. (A) Principal Component Analysis (PCA) of each point mutation, truncation, insertion and first-determined WT Asyn fibril protomer, showing the PC2 vs PC1 plot. DBSCAN was used to cluster the points in PC space, which are labeled by their four-character PDB identifier code. Points and codes in blue were found to be part of a cluster, which corresponds to Cluster 1 as shown previously^4^. (B) Per-residue root-mean-square fluctuation (RMSF) plot for each protomer in the group, shown between residues V40-K96. (C) Alignment of clustered protomers shows characteristic Class 1 Greek key fold. (D) Conserved binding pocket observed among all Cluster 1 fibrils, with the sidechains of K43, K45, V48 and H50 exposed to the solvent and amenable to ligand binding.

## DISCUSSION

We produced a large quantity of the hereditary mutant A30P Asyn fibrils to enable their high-resolution structure determination. Several thousand SSNMR distance restraints and dihedral angles were used to generate a structural model based on Xplor-NIH simulated annealing calculations. The fibril fold is unsurprisingly similar to that of *in vitro* wild-type fibrils, other *in vitro* hereditary point mutation fibrils, as well as *ex vivo* MSA and Juvenile Onset Synucleinopathy (JOS) fibrils, all defined by an orthogonal Greek key architecture and druggable pocket of exposed sidechains between K43-H50. These results not only expand the library of high-resolution Asyn fibril structures but suggest that the Greek key fold is accessible across a wide variety of constructs and preparation conditions.

We have also achieved the determination of fibril structure to atomic-level resolution using only one u-^13^C, ^15^N labelled SSNMR sample. Previously, amyloid fibril structure determination to a similar resolution by SSNMR required several samples with different isotopic labelling schemes, alongside complex experiments to calculate specific heteronuclear distance restraints^35, 47, 52, 99^. Using only a combination of five 2D and 3D homonuclear ^13^C correlation spectra with varying DARR mixing times, we were able to generate thousands of distance restraints which were then used to construct a high-resolution model of A30P Asyn fibrils. While our SSNMR sample contained approximately 16 mg of material packed into a 3.2 mm rotor, it should be possible to apply the same approaches with deuterated samples, smaller sample quantity, smaller rotor size, higher-field magnets and faster magic angle spinning to increase the resolution while maintaining similar sensitivity. Ongoing efforts by our group and others in the field are already starting to show promise in this direction^100-103^.

Since structurally similar fibrils will have similar backbone chemical shifts and cross peaks in their SSNMR spectra, we were able to quantitatively compare the spectra of several Asyn fibril polymorphs and identify Class 1A fibrils via clustering pairwise ZNCC scores^4, 104^. While we were able to use this quantitative tool to determine structural similarity, it also has the promise of being able to rapidly classify fibril structures from new samples via a single SSNMR spectrum. For sample quantities on the order of a few milligrams, high-sensitivity and well-resolved 2D ^15^N-^13^Ca spectra can be collected in under 24 hours (this study). Similar quality can be achieved for samples of smaller quantity by signal averaging for longer, which is especially important for fibril samples amplified from diseased patient tissue or CSF, where starting material is on the level of picograms to nanograms and faithful amplification is still on the order of a few milligrams^47, 105, 106^. Given that structures have only been determined for a handful of patient-derived fibrils, this structural classification procedure could be employed to rapidly classify the structures of patient-derived fibrils not only from synucleinopathy patients, but patients with tauopathies, TDP43-opathies and other diseases marked by fibril formation such as Alzheimer Disease, Huntington’s Disease, and Type II diabetes. Though cryo-EM is emerging as the preferred method in the field for determining structures of fibrils, this SSNMR-based approach is advantageous for testing *in vitro* conditions which might recapitulate disease-relevant folds. Additionally, using both cryo-EM and SSNMR to determine structure can be complementary, as a means of validation or by filling in gaps in a structure that one method alone would not be able to determine.

While the structure of *in vitro*-formed A30P Asyn fibrils adopts a Greek key topology and falls into Class 1A of fibrils^4^, it is likely that the structure of A30P Asyn fibrils *in vivo* is quite different and resembles the Lewy fold found in both *ex vivo* and *ex vivo* seeded fibrils from patients with PD, DLB and PDD^39, 47^. A recent preprint by Yang et al. describes the structures of *ex vivo* G51D and H50Q patient-derived Asyn fibrils, revealing a Lewy fold topology with a characteristic β-arc between G51-G67 and an extended β-arc between G73-A91^107^. While no structure of *ex vivo* A30P Asyn fibrils has been reported to this date, it is likely they will also exhibit the Lewy Fold.

That *in vitro* A30P Asyn fibrils display a Greek key fold is not surprising. Under the formation conditions of 50 mM sodium phosphate, pH 7.4 and 200 rpm shaking for a period of 6 weeks, there is a high likelihood that the fibrils will be mature, largely homogenous and adapt a thermodynamically favorable fold. A recent preprint shows that both WT and A53T Asyn are capable of forming fibrils with different folds even when using identical formation conditions, suggesting that fibrils form stochastically^108^. Given that the fibrils formed in this preprint were subjected to only 10-14 days incubation, it is likely that at this stage the fibrils have yet to find the global minimum on their formation energy landscape. In our data for both A30P fibrils (this study) and WT fibrils^35^ – both of which used incubation periods of at least 21 days – only one set of peaks is present, which is indicative of a single population of fibrils. While it is possible that incubating A30P Asyn for a smaller period of time might result in different or multiple folds, the Greek key fold is likely the thermodynamically favorable fold.

The ability for structures from *in vitro* mutants A30P, H50Q, A53T, A53E, 1-100, 1-121 and the MAAEKT-insertion fibrils to adapt a Greek key fold suggests that mutation does not drive structure but can instead restrict the fibril formation energy landscape and increase the probability of contacts forming which induce such a fold. E46K fibrils severely restrict this landscape by limiting the formation of the E46-K80 salt bridge, a key stability-contributing contact in Cluster 1 fibrils. While the other hereditary mutations introduce charge, polarity or conformational flexibility, they do not disrupt key contacts which stabilize the fold of Cluster 1 fibrils. Given the new structures of H50Q and G51D derived from PD patient tissue adapt the Lewy Fold at the protomer level, it is likely that the mutations themselves disrupt the normal physiological processes Asyn is involved in and increase the propensity of aggregation, leading to a common fold rather than mutation-specific folds.

## ASSOCIATED CONTENT

### Supporting Information

This material is available free of charge via the Internet at http://pubs.acs.org.

- Extent of SSNMR backbone and sidechain assignments and secondary structure of A30P fibrils, Random Coil Index S^2^ order parameter by residue, as well as 1D ^13^C CP and DP spectra showing rigid residues and length of fibril core; Protein backbone walk using 3D ^15^N-^13^Cα-^13^CX, 3D ^13^Cα-^15^N-(^13^C’)-^13^CX and 3D ^15^N-^13^C’-^13^CX spectra; ^13^C and ^15^N Chemical shift differences between previous A30P sample (BMRB: 17648) and this sample; Series of 1D ^15^N CP spectra with varying ^1^H-^15^N CP contact times and 1D ^15^N DP, as well as 2D ^15^N-^13^Cα spectra collected at 0° C and -50° C to show P30’s location outside of the fibril core; Strips from 2D ^13^C-^13^C with 500 ms DARR mixing and 2D ^13^C-^13^C with 125 ms DARR mixing spectra to show unambiguous long-range distances; Overlay of experimental and NMRFAM-BPHON simulated 2D ^15^N-^13^Cα and 2D ^13^C-^13^C with 500 ms DARR mixing spectra; Simulated 1D ^13^C CP and DP spectra with varying sequence lengths, as well as SSD-NMR predicted secondary structure element percentages; Table of SSNMR experiments with acquisition parameters, experiment time, and initial and final distance restraints derived from their peak lists; NMR resonance list with backbone and sidechain chemical shifts for residues between T33 and D98; Backbone dihedral angles predicted with TALOS-N; χ1 angles and rotamer states predicted with TALOS-N; Unambiguous manual distance restraints used as a standalone energy term in Xplor-NIH structure calculations; Summary of simulated annealing protocols in the PASD algorithm; List of Xplor-NIH energy terms used.

## Author Contributions

Conceptualization: M.H.M. and C.M.R. Methodology: M.H.M and C.M.R. Software: M.H.M., O.A.W., C.D.S., and C.M.R. Validation: M.H.M., O.A.W., C.G.B., and C.M.R. Formal Analysis:M.H.M. Investigation: M.H.M, B.D.H., R.H., and C.M.R. Resources: C.M.R. Data curation: M.H.M. Writing – Original draft: M.H.M. Writing – Reviewing and editing: M.H.M., O.A.W., C.G.B., R.H., D.D.D., P.T.K., E.R.W., and C.M.R. Visualization: M.H.M. Funding acquisition: P.T.K., E.R.W., and C.M.R.

## Notes

Any additional relevant notes should be placed here.

## ACKNOWLEDGMENTS

This work was supported by 2RF1NS110436-06 (NINDS), P41GM136463 (NIGMS), the William H. Peterson Graduate Fellowship in Biochemistry (awarded to M.H.M. and B.D.H.), the Molecular Biophysics Predoctoral Training Grant T32GM130550 (NIGMS) (awarded to O.A.W.), and the NIH Ruth L. Kirschstein Fellowship F32GM149118 (NIGMS) (awarded to C.G.B.). C.D.S. was supported by the Intramural Research Program of the National Institute of Diabetes and Digestive and Kidney Diseases, National Institutes of Health. We would like to thank Dr. Gemma Comellas and Dr. Luisel Lemkau for their previous work on A30P fibrils. We would also like to thank Dr. Rajat Garg and Barry DeZonia for their continued assistance with and maintenance of the NMRFAM-BPHON program and workflows used in this manuscript.

---

Authors are required to submit a graphic entry for the Table of Contents (TOC) that, in conjunction with the manuscript title, should give the reader a representative idea of one of the following: A key structure, reaction, equation, concept, or theorem, etc., that is discussed in the manuscript. Consult the journal’s Instructions for Authors for TOC graphic specifications.

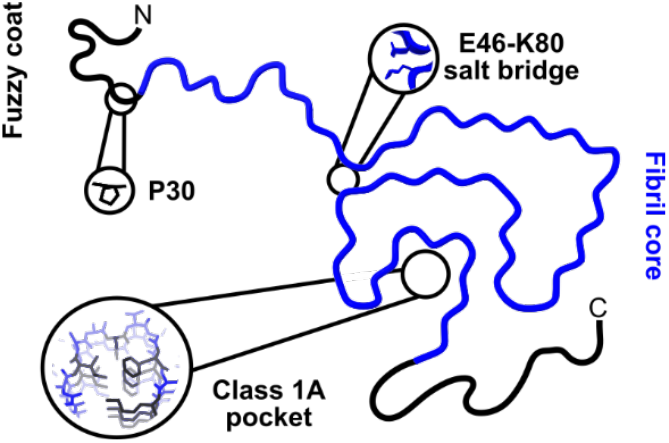

