## Supporting Information for "*In vitro*-prepared A30P alpha-synuclein fibrils adopt the conserved and disease-relevant Greek key fold"

^1^Department of Biochemistry, University of Wisconsin-Madison, Madison, WI, 53706, USA; ^2^Graduate Program in Biophysics, University of Wisconsin-Madison, Madison, WI, 53706, USA; ^3^Department of Chemistry, University of Wisconsin-Madison, Madison, WI, 53706, USA; ^4^Department of Neurology and Hope Center for Neurological Disorders, Washington University School of Medicine, St. Louis, Missouri, 63110, USA; ^5^Cryo-Electron Microscopy Research Center, University of Wisconsin-Madison, Madison, WI, 53706, USA; ^6^Midwest Center for Cryo-Electron Tomography, Department of Biochemistry, University of Wisconsin-Madison, Madison, WI, 53706, USA; ^7^Morgridge Institute for Research, Madison, WI, 53715, USA; ^8^Imaging Sciences Laboratory, Center for Information Technology, National Institutes of Health, Bethesda, MD, 20817, USA; ^9^National Magnetic Resonance Facility at Madison, University of Wisconsin-Madison, Madison, WI, 53706, USA

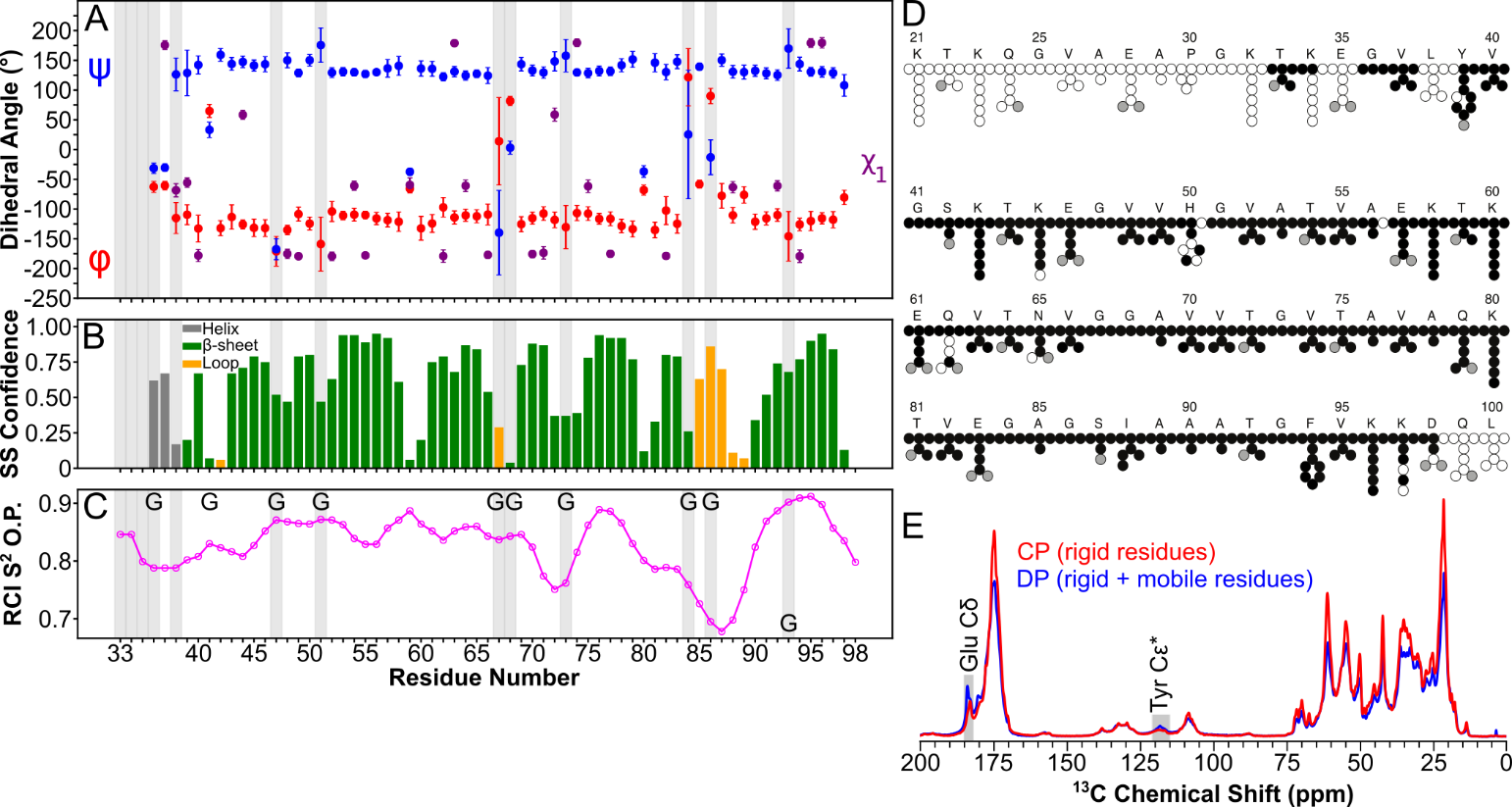

**Figure S1:** SSNMR assignments and secondary structure. (A) Backbone and χ1 dihedral angles of A30P Asyn fibrils predicted with the program TALOS-N from backbone and sidechain chemical shift assignments^1^. ψ (blue), φ (red) and χ1 (purple) dihedral angles plotted on an absolute degree scale, showing error bars in some cases spanning several tens of degrees, mostly in cases where confidence is low. (B) TALOS-N prediction confidence for secondary structure elements, with residues predicted as helices in grey, β-sheets in green and loops in orange. (C) RCI S^2^ order parameter for A30P Asyn fibrils, detailing relative degree of rigidity for regions within the fibril core. Score predicted for residues 33-98 even though dihedral angles could not be predicted for 33-35 due to lack of backbone chemical shifts. Glycine residues marked by a G, most of which were left out of the Xplor-NIH dihedral angle energy term. Residues shaded in grey were labeled “Warn” by TALOS-N and reflect residues with poor confidence in dihedral angle prediction. (D) Extent of ^13^C and ^15^N chemical shift assignments for each residue within the fibril core, shown as a tree diagram. (E) Overlay of 1D ^13^C cross polarization spectrum (rigid residues; fibril core) with 1D ^13^C direct polarization spectrum (rigid + mobile residues; fibril core + fuzzy coat). Note the increase in the peak intensity within the DP spectrum at ~183 ppm and ~117 ppm (shaded in grey), which are specific for glutamic acid Cδ resonances and tyrosine Cε* resonances, respectively, which are highly abundant within the dynamically disordered C-terminus.

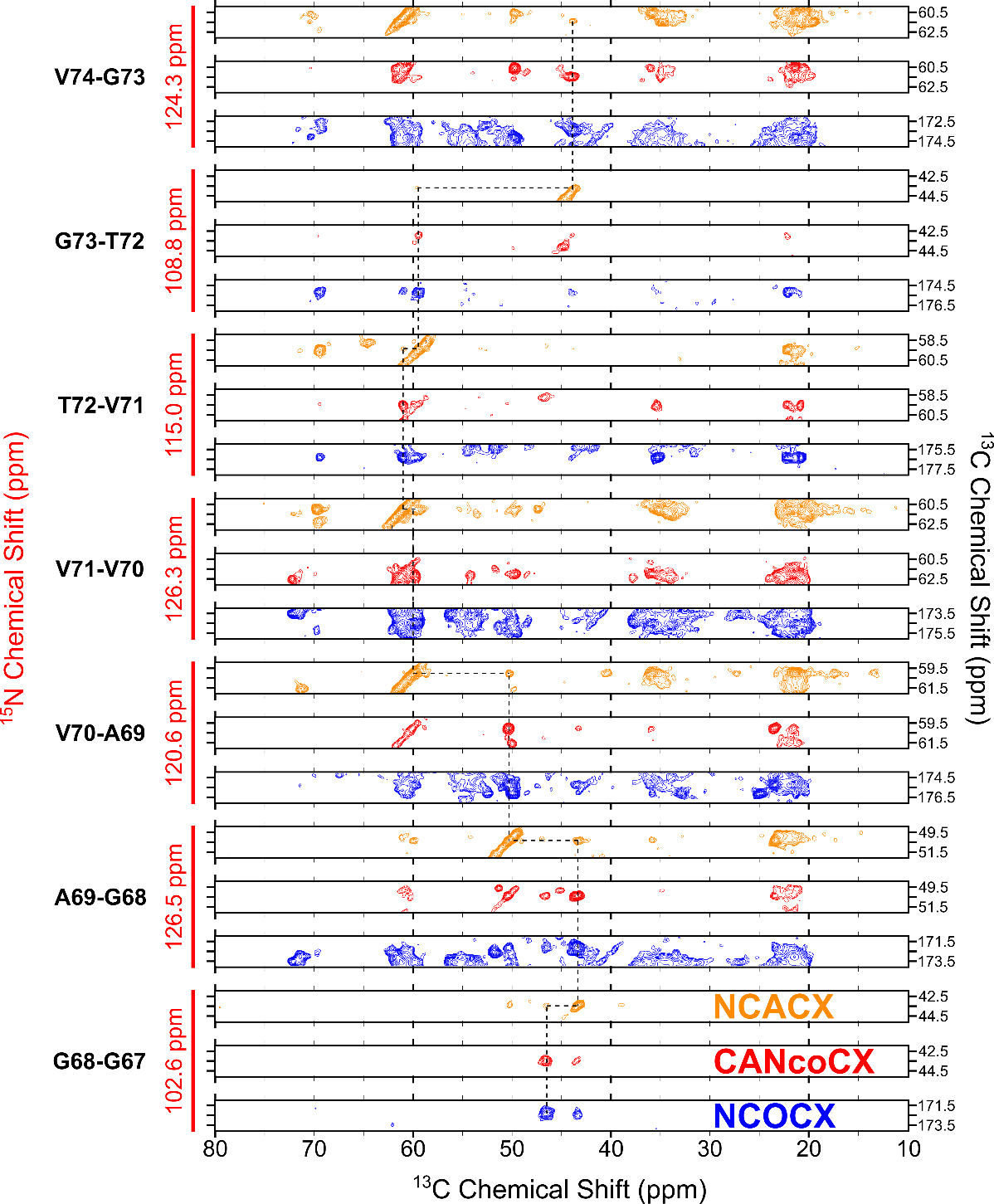

**Figure S2:** Protein backbone walk to assign resonances of A30P fibrils. Example stretch between G67-V74, connecting backbone Cα, N and C’ resonances between sequential residues. 3D ^15^N-^13^Cα-^13^CX (orange; labeled as NCACX), ^13^Cα-^15^N-(^13^C’)-^13^CX (red; labeled as CANcoCX) and ^15^N-^13^C’-^13^CX (blue; labeled as NCOCX) spectra were used to establish backbone connectivity. Experimental parameters for these spectra can be found in **Table S1**.

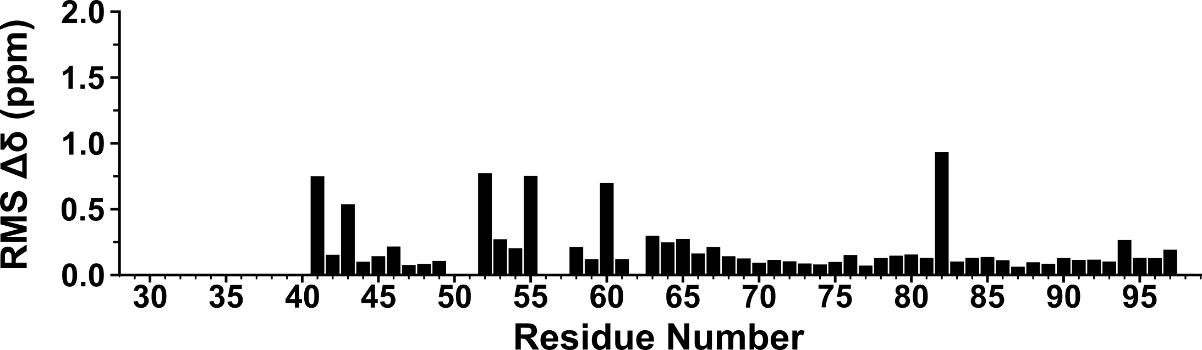

**Figure S3:** ^13^C and ^15^N Chemical Shift Differences (CSDs) between this sample (BMRB: 31269) and A30P Asyn fibrils deposited under BMRB: 17648^2^.

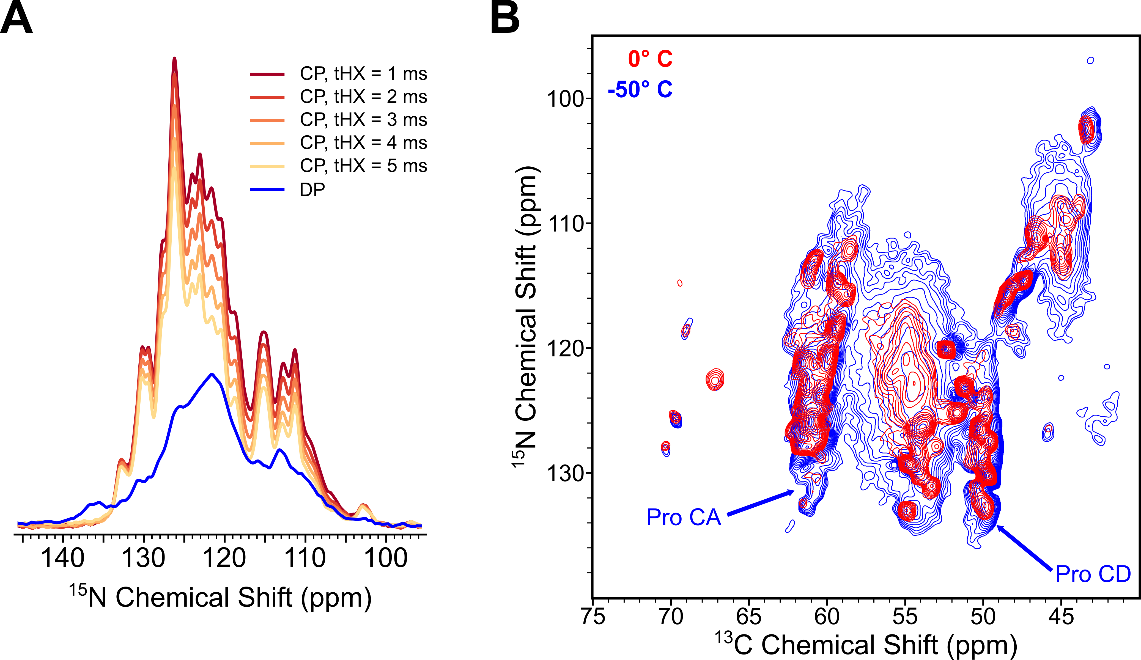

**Figure S4:** P30 is located outside of fibril core. (A) Series of 1D ^15^N spectra with increasing ^1^H-^15^N cross-polarization contact time, along with 1D ^15^N direct polarization spectrum (rigid + disordered residues). (B) Overlay of 2D ^15^N-^13^Cα spectra collected at VT set points of 0° C (red) and -50° C (blue). Terminal molecular motions freeze out at -50° C, resulting in broadening of peaks in every region of spectrum as well as appearance of new peaks in both Proline N-Cα and N-Cδ regions.

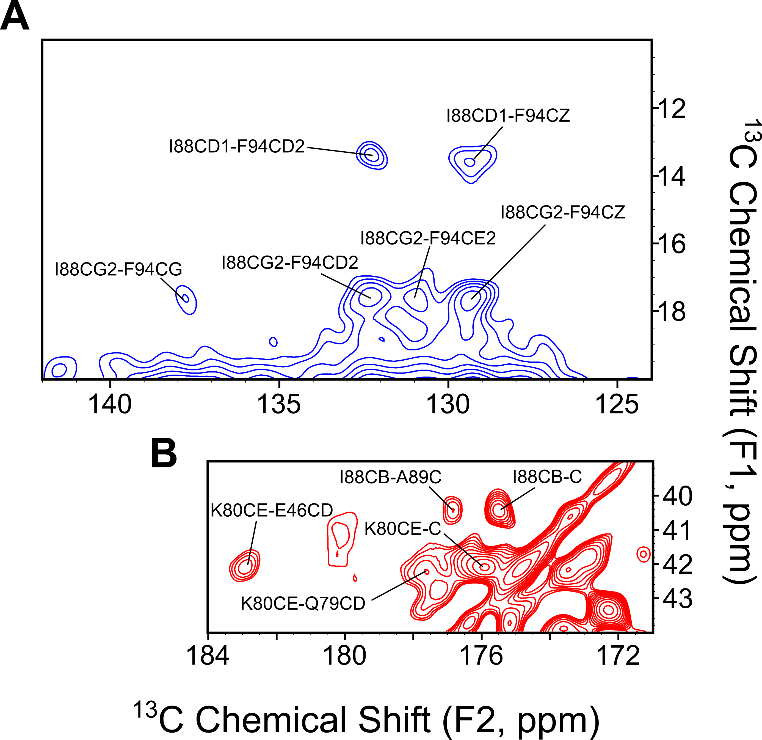

**Figure S5:** (A) Region of 2D ^13^C-^13^C with 500 ms DARR mixing spectrum showing sidechains of I88 and F94 in close proximity, forming the basis of the hydrophobic pocket characteristic of a closed-in Greek key. (B) Region of 2D ^13^C-^13^C with 125 ms DARR mixing spectrum showing interaction between K80Cε and E46Cδ at a distance of ≤ 8 Å.

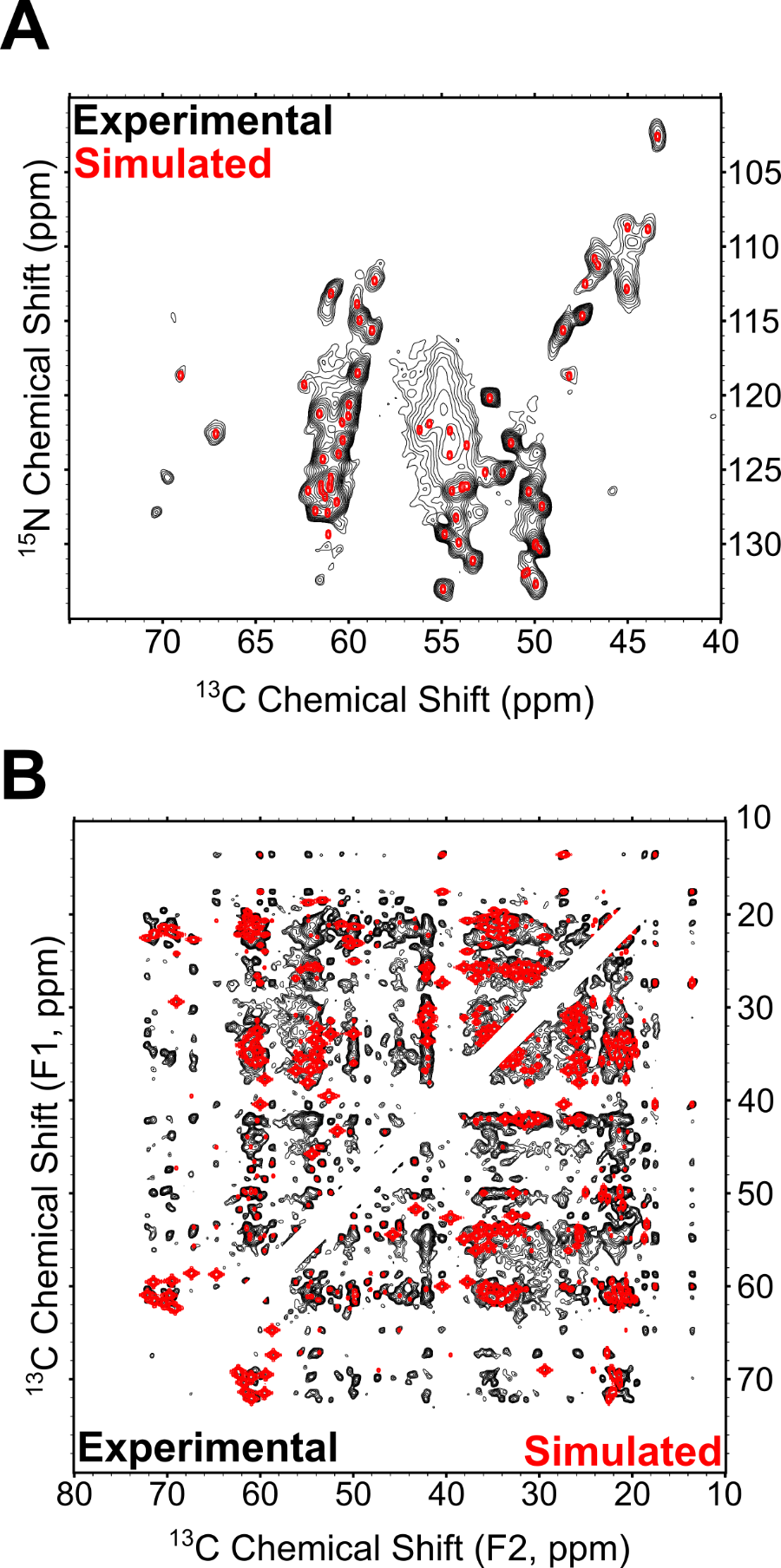

**Figure S6:** Overlay of experimental spectra with spectra simulated using NMRFAM-BPHON and experimental resonance list (BMRB: 31269). (A) 2D ^15^N-^13^Cα spectra, experimental in black and simulated in red. ZNCC score is 0.48. (B) 2D ^13^C-^13^C spectra, experimental is with 500 ms DARR mixing (black) and simulated is with 200 ms DARR mixing to achieve atomic distances of 10 Å (red). ZNCC score is 0.31.

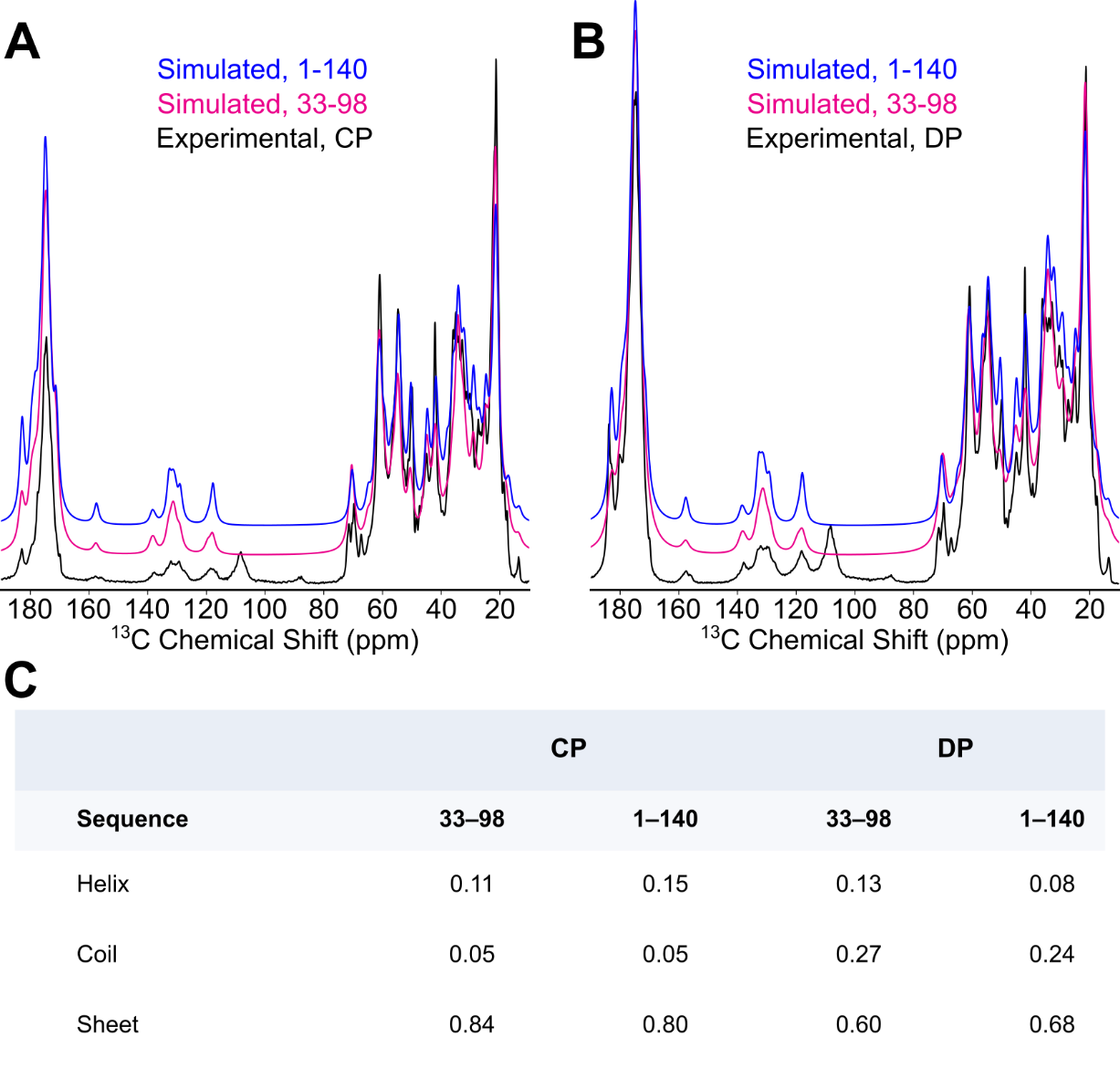

**Figure S7**: Spectral simulation and secondary structure element prediction using SSD-NMR. (A) Overlay of experimental ^13^C 1D CP spectrum (black), simulated ^13^C 1D under the assumption of rigid residues spanning T33-D98 (magenta), and simulated ^13^C 1D under the assumption of rigid residues spanning M1-A140 (blue). (B) Overlay of experimental ^13^C 1D DP spectrum (black), simulated ^13^C 1D assuming the signals arise from T33-D98, and simulated ^13^C 1D assuming the signals arise from M1-A140. (C) Secondary structure element predictions for each spectrum, based on their similarity to the experimental spectra. Carbonyl sideband peaks at roughly 110 ppm in the experimental spectra are not found in the simulated spectra.

**Table S1:** Table of SSNMR Experiments^a^.

| Experiment number | Experiment name | NMR Parameters | Time (min) | Peak list used for PASD | | Initial distance restraints | | Final distance restraints |
| --- | --- | --- | --- | --- | --- | --- | --- | --- |
| 1 | 1D ^13^C CP | nt = 32, τ_rd_ (d1) = 1.5 s, τ_acq_ = 10 ms, ν_1H_ = 113.6 kHz, ν_13C_ = 113.6 kHz, τ_HC_ = 900 µs, ν_1H, SPINAL_= 82.4 kHz | 1 | No | - | | - | |
| 2 | 1D ^13^C DP | nt = 128, τ_rd_ (d1) = 5 s, τ_acq_ = 30 ms, ν_1H_ = 92.6 kHz, ν_13C_ = 125.0 kHz, ν_1H, SPINAL_= 84.9 kHz | 12 | No | - | | - | |
| 3 | 1D ^15^N CP | nt = 256, τ_rd_ (d1) = 1.5 s, τ_acq_ = 30 ms, ν_1H_ = 108.7 kHz, ν_15N_ = 52.1 kHz, τ_HC_ = 1000, 2000, 3000, 4000 and 5000 µs, ν_1H, SPINAL_= 76.1 kHz | 33 | No | - | | - | |
| 4 | 1D ^15^N DP | nt = 2048, τ_rd_ (d1) = 30 s, τ_acq_ = 40 ms, ν_1H_ = 108.7 kHz, ν_15N_ = 50.0 kHz, ν_1H, SPINAL_= 76.1 kHz | 918 | No | - | | - | |
| 5 | 2D ^13^C-^13^C | nt = 4, τ_rd_ (d1) = 1.5 s, τ_acq_ = 15 ms, ν_1H_ = 113.6 kHz, ν_13C_ = 108.7 kHz, τ_HC_ = 900 µs, ν_1H, SPINAL_= 82.4 kHz, τ_DARR_= 50 ms | 108 | Yes | 354 | | 276 | |
| 6 | 2D ^13^C-^13^C | nt = 4, τ_rd_ (d1) = 1.5 s, τ_acq_ = 15 ms, ν_1H_ = 113.6 kHz, ν_13C_ = 108.7 kHz, τ_HC_ = 900 µs, ν_1H, SPINAL_= 82.4 kHz, τ_DARR_= 125 ms | 1053 | Yes | 604 | | 427 | |
| 7 | 2D ^13^C-^13^C | nt = 8, τ_rd_ (d1) = 1.5 s, τ_acq_ = 18 ms, ν_1H_ = 92.6 kHz, ν_13C_ = 125.0 kHz, τ_HC_ = 600 µs, ν_1H, SPINAL_= 84.9 kHz, τ_DARR_= 500 ms | 2184 | Yes | 790 | | 604 | |
| 8 | 2D ^15^N-^13^CA | nt = 16, τ_rd_ (d1) = 1.5 s, τ_acq_ = 18 ms, ν_1H_ = 113.6 kHz, ν_13C_ = 108.7 kHz, ν_15N_ = 52.1 kHz, τ_HN_ = 1800 µs, τ_NCa_ = 6000 µs, ν_1H, SPINAL_= 82.4 kHz | 420 | No | - | | - | |
| 9 | 2D ^15^N-^13^CA | nt = 8, τ_rd_ (d1) = 1.5 s, τ_acq_ = 18 ms, T = 223.15 K, ν_1H_ = 113.6 kHz, ν_13C_ = 108.7 kHz, ν_15N_ = 52.1 kHz, τ_HN_ = 4500 µs, τ_NCa_ = 5000 µs, ν_1H, SPINAL_= 79.6 kHz | 1802 | No | - | | - | |
| 10 | 2D ^15^N-^13^ca-^13^CX | nt = 16, τ_rd_ (d1) = 1.5 s, τ_acq_ = 18 ms, ν_1H_ = 113.6 kHz, ν_13C_ = 108.7 kHz, ν_15N_ = 52.1 kHz, τ_HN_ = 1800 µs, τ_NCa_ = 6000 µs, ν_1H, SPINAL_= 82.4 kHz, τ_DARR_= 50 ms | 1360 | No | - | | - | |
| 11 | 3D ^15^N-^13^CA-^13^CX | nt = 4, τ_rd_ (d1) = 1.5 s, τ_acq_ = 18 ms, ν_1H_ = 113.6 kHz, ν_13C_ = 108.7 kHz, ν_15N_ = 52.1 kHz, τ_HN_ = 1800 µs, τ_NCa_ = 6000 µs, ν_1H, SPINAL_= 82.4 kHz, τ_DARR_= 50 ms | 4368 | No | - | | - | |
| 12 | 3D ^15^N-^13^CO-^13^CX | nt = 4, τ_rd_ (d1) = 1.5 s, τ_acq_ = 18 ms, ν_1H_ = 113.6 kHz, ν_13C_ = 108.7 kHz, ν_15N_ = 52.1 kHz, τ_HN_ = 1800 µs, τ_NCO_ = 6000 µs, ν_1H, SPINAL_= 82.4 kHz, τ_DARR_= 50 ms | 5472 | No | - | | - | |
| 13 | 3D ^13^CA-^15^N-^13^CO | nt = 8, τ_rd_ (d1) = 1.5 s, τ_acq_ = 18 ms, ν_1H_ = 113.6 kHz, ν_13C_ = 104.2 kHz, ν_15N_ = 52.1 kHz, τ_HC_ = 550 µs, τ_CaN_ = 5000 µs, τ_NCO_ = 5000 µs, ν_1H, SPINAL_= 82.4 kHz | 3383 | No | - | | - | |
| 14 | 3D ^13^CA-^15^N-^13^co-^13^CX | nt = 4, τ_rd_ (d1) = 1.5 s, τ_acq_ = 18 ms, ν_1H_ = 113.6 kHz, ν_13C_ = 104.2 kHz, ν_15N_ = 52.1 kHz, τ_HC_ = 560 µs, τ_CaN_ = 6000 µs, τ_NCO_ =6000 µs, ν_1H, SPINAL_= 82.4 kHz, τ_DARR_= 50 ms | 11090 | No | - | | - | |
| 15 | 3D ^13^C-^13^C-^13^C | nt = 16, τ_rd_ (d1) = 1.25 s, τ_acq_ = 15 ms, ν_1H_ = 104.2 kHz, ν_13C_ = 83.3 kHz, τ_HC_ = 1600 µs, ν_1H, SPINAL_= 84.6 kHz, τ_DARR, 1_= 50 ms, τ_DARR, 2_= 500 ms | 21558 | Yes | 1894 | | 963 | |
| 16 | 3D ^13^C-^13^C-^13^C | nt = 2, τ_rd_ (d1) = 1.5 s, τ_acq_ = 18 ms, ν_1H_ = 92.6 kHz, ν_13C_ = 125.0 kHz, τ_HC_ = 600 µs, ν_1H, SPINAL_= 84.9 kHz, τ_DARR, 1_= 50 ms, τ_DARR, 2_= 500 ms | 10246 | Yes | 4316 | | 1917 | |

^a^All experiments were collected at a VT set point temperature of 273.15 K (unless noted otherwise), *ν_MAS_* = 12.5 kHz, B_0_ = 750 MHz on a Varian VNMRS spectrometer running Open VNMRJ 3.1, with a Balun 3.2 mm probe. Symbols: nt = number of scans, τ_rd_ = recycle delay between scans, τ_acq_= acquisition time, ν_1H_ = ^1^H radiofrequency (RF) field strength, ν_13C_ = ^13^C RF field strength, ν_15N_ = ^15^N RF field strength, τ_HC_ = cross-polarization contact time from ^1^H to ^13^C channel, τ_CaN_ = Contact time from ^13^Ca channel to ^15^N channel, τ_NCO_ = Contact time from ^15^N channel to ^13^Co channel, ν_1H, SPINAL_= ^1^H RF field strength during SPINAL decoupling, τ_DARR_= DARR mixing time.

**Table S2:** Backbone and Sidechain Resonance Assignments^b^.

| Residues | N | Nζ | C’ | Cα | Cβ | Cγ/γ_1_ | Cγ_2_ | Cδ/δ_1_ | Cδ_2_ | Cε/ε_1_ | Cε_2_ | Cζ |
| --- | --- | --- | --- | --- | --- | --- | --- | --- | --- | --- | --- | --- |
| T33 | 119.3 |  | 172.2 | 62.4 | 69.2 |  | 22.2 |  |  |  |  |  |
| K34 | 131.9 |  |  | 50.4 |  |  |  |  |  |  |  |  |
| G36 | 112.5 |  | 174.1 | 47.3 |  |  |  |  |  |  |  |  |
| V37 | 118.7 |  | 177.8 | 69.0 | 29.4 | 24.2 | 22.4 |  |  |  |  |  |
| Y39 |  |  | 174.6 | 52.6 | 39.5 | 127.2 |  | 128.7 |  | 120.1 | 118.8 | 156.0 |
| V40 | 121.8 |  | 177.2 | 60.3 | 34.6 | 22.0 | 21.2 |  |  |  |  |  |
| G41 | 118.7 |  | 174.2 | 48.1 |  |  |  |  |  |  |  |  |
| S42 | 112.3 |  | 171.2 | 58.6 | 67.4 |  |  |  |  |  |  |  |
| K43 | 122.4 |  | 175.6 | 54.6 | 36.8 | 25.7 |  | 31.6 |  | 42.5 |  |  |
| T44 | 113.9 |  | 175.2 | 59.5 | 71.5 |  | 22.3 |  |  |  |  |  |
| K45 | 122.3 |  | 173.6 | 56.2 | 36.8 | 26.9 |  | 30.8 |  | 42.2 |  |  |
| E46 | 126.2 |  | 174.8 | 53.9 | 32.3 | 34.8 |  | 182.9 |  |  |  |  |
| G47 | 115.6 |  | 172.6 | 48.5 |  |  |  |  |  |  |  |  |
| V48 | 118.5 |  | 174.5 | 59.5 | 37.7 | 24.0 | 20.7 |  |  |  |  |  |
| V49 | 126.4 |  | 173.4 | 61.5 | 34.1 | 22.4 | 20.3 |  |  |  |  |  |
| H50 | 126.1 |  |  | 53.9 | 34.7 |  |  |  | 116.4 | 137.7 |  |  |
| G51 | 108.7 |  | 170.7 | 45.0 |  |  |  |  |  |  |  |  |
| V52 | 126.1 |  | 174.6 | 61.0 | 33.4 | 22.8 | 20.8 |  |  |  |  |  |
| A53 | 132.7 |  | 175.4 | 49.9 | 21.3 |  |  |  |  |  |  |  |
| T54 | 121.2 |  | 173.1 | 61.6 | 71.4 |  | 21.9 |  |  |  |  |  |
| V55 | 126.8 |  | 175.6 | 61.3 | 33.8 | 21.8 | 19.8 |  |  |  |  |  |
| A56 | 132.0 |  |  | 50.6 | 22.6 |  |  |  |  |  |  |  |
| E57 | 130.1 |  | 175.2 | 50.0 | 32.8 | 36.0 |  | 183.6 |  |  |  |  |
| K58 | 123.4 | 34.1 | 176.3 | 53.7 | 36.3 | 25.8 |  | 30.1 |  | 41.9 |  |  |
| T59 | 122.6 |  | 174.1 | 67.1 | 67.2 |  | 22.7 |  |  |  |  |  |
| K60 | 121.9 | 34.2 | 173.3 | 55.7 | 35.5 | 26.0 |  | 33.7 |  | 42.0 |  |  |
| E61 | 128.2 |  | 174.6 | 54.2 | 33.5 | 35.2 |  | 182.7 |  |  |  |  |
| Q62 | 124.0 |  | 175.1 | 54.6 |  |  |  | 179.5 |  |  |  |  |
| V63 | 129.4 |  | 174.8 | 61.1 | 35.3 | 21.0 | 20.3 |  |  |  |  |  |
| T64 | 126.4 |  | 172.5 | 62.2 | 69.6 |  | 21.4 |  |  |  |  |  |
| N65 | 125.2 |  | 172.6 | 51.7 | 43.3 | 174.7 |  |  |  |  |  |  |
| V66 | 127.2 |  | 177.7 | 60.6 | 33.1 | 21.2 | 20.5 |  |  |  |  |  |
| G67 | 111.3 |  | 172.3 | 46.6 |  |  |  |  |  |  |  |  |
| G68 | 102.6 |  | 172.2 | 43.4 |  |  |  |  |  |  |  |  |
| A69 | 126.5 |  | 175.2 | 50.3 | 23.4 |  |  |  |  |  |  |  |
| V70 | 120.6 |  | 174.5 | 60.0 | 35.7 | 23.3 | 21.4 |  |  |  |  |  |
| V71 | 126.2 |  | 176.3 | 61.0 | 35.2 | 22.1 | 20.8 |  |  |  |  |  |
| T72 | 115.0 |  | 175.1 | 59.4 | 69.5 |  | 22.1 |  |  |  |  |  |
| G73 | 108.8 |  | 173.4 | 43.9 |  |  |  |  |  |  |  |  |
| V74 | 124.3 |  | 175.0 | 61.4 | 34.9 | 21.1 | 19.6 |  |  |  |  |  |
| T75 | 127.8 |  | 171.9 | 61.8 | 70.4 |  | 21.3 |  |  |  |  |  |
| A76 | 130.3 |  | 174.2 | 49.7 | 21.3 |  |  |  |  |  |  |  |
| V77 | 124.0 |  | 172.8 | 60.5 | 35.9 | 21.3 | 20.4 |  |  |  |  |  |
| A78 | 130.0 |  | 176.2 | 49.9 | 25.0 |  |  |  |  |  |  |  |
| Q79 | 120.2 |  | 176.3 | 52.4 | 32.9 | 31.4 |  | 177.6 |  |  |  |  |
| K80 | 123.0 | 33.6 | 175.9 | 60.3 | 32.5 | 26.6 |  | 31.4 |  | 42.2 |  |  |
| T81 | 113.2 |  | 173.5 | 60.9 | 72.1 |  | 22.5 |  |  |  |  |  |
| V82 | 126.0 |  | 174.7 | 61.5 | 34.1 | 20.7 | 20.1 |  |  |  |  |  |
| E83 | 126.1 |  | 174.9 | 53.6 | 33.9 | 36.0 |  | 183.0 |  |  |  |  |
| G84 | 112.8 |  | 173.4 | 45.0 |  |  |  |  |  |  |  |  |
| A85 | 131.1 |  | 179.1 | 53.3 | 18.4 |  |  |  |  |  |  |  |
| G86 | 110.8 |  | 173.6 | 46.8 |  |  |  |  |  |  |  |  |
| S87 | 115.6 |  | 173.4 | 58.7 | 64.7 |  |  |  |  |  |  |  |
| I88 | 121.4 |  | 175.5 | 60.0 | 40.4 | 27.4 | 17.6 | 13.6 |  |  |  |  |
| A89 | 129.4 |  | 176.8 | 54.8 | 18.7 |  |  |  |  |  |  |  |
| A90 | 123.2 |  | 174.5 | 51.3 | 21.0 |  |  |  |  |  |  |  |
| A91 | 127.5 |  | 175.4 | 49.6 | 23.1 |  |  |  |  |  |  |  |
| T92 | 125.6 |  | 174.5 | 61.0 | 69.8 |  | 21.9 |  |  |  |  |  |
| G93 | 114.7 |  | 169.9 | 47.4 |  |  |  |  |  |  |  |  |
| F94 | 126.4 |  | 173.6 | 54.4 | 45.8 | 137.9 |  | 132.2 | 132.2 | 131.1 | 130.7 | 129.3 |
| V95 | 127.9 |  | 171.3 | 61.1 | 34.8 | 22.2 | 20.5 |  |  |  |  |  |
| K96 | 133.0 | 29.3 | 173.0 | 54.9 | 38.1 | 25.7 |  | 31.1 |  | 41.8 |  |  |
| K97 | 129.9 |  | 175.0 | 54.1 | 35.4 | 25.4 |  | 32.2 |  | 41.9 |  |  |
| D98 | 125.2 |  |  | 52.7 | 39.5 |  |  |  |  |  |  |  |

^b^Assigned resonances of A30P fibrils used for distance restraint assignment in PASD ^3^. Resonance frequency values listed in units of parts-per-million (ppm). Underlined resonance frequencies were not able to be assigned using the data but given the average value found in the BMRB and subjected to variable tolerances in the initMatch step of PASD.

**Table S3:** Backbone dihedral angles predicted using TALOS-N^c^

| Residue | Phi (φ) | Δφ | Psi (ψ) | Δψ | TALOS-N rank | |
| --- | --- | --- | --- | --- | --- | --- |
| G36 | -62.680 | 9.231 | -31.360 | 8.586 | | Warn |
| V37 | -60.548 | 7.220 | -30.438 | 6.330 | | Strong |
| L38 | -115.132 | 25.942 | 126.273 | 27.606 | | Warn |
| Y39 | -109.538 | 16.567 | 128.840 | 38.415 | | Strong |
| V40 | -132.730 | 22.460 | 142.295 | 15.213 | | Strong |
| G41 | 64.810 | 11.011 | 33.142 | 13.004 | | Strong |
| S42 | -132.093 | 12.504 | 159.181 | 10.721 | | Strong |
| K43 | -113.445 | 20.398 | 143.874 | 10.755 | | Strong |
| T44 | -125.837 | 7.827 | 147.332 | 9.739 | | Strong |
| K45 | -131.643 | 14.682 | 141.269 | 9.400 | | Strong |
| E46 | -132.018 | 14.223 | 143.531 | 12.779 | | Strong |
| G47 | -171.059 | 25.025 | -167.544 | 17.710 | | Warn |
| V48 | -135.118 | 7.922 | 150.111 | 12.672 | | Strong |
| V49 | -108.531 | 11.717 | 128.873 | 6.542 | | Strong |
| H50 | -124.403 | 10.818 | 150.199 | 9.757 | | Strong |
| G51 | -158.892 | 45.329 | 175.603 | 28.771 | | Warn |
| V52 | -104.103 | 16.934 | 129.301 | 8.158 | | Strong |
| A53 | -110.997 | 7.232 | 130.958 | 7.089 | | Strong |
| T54 | -109.405 | 10.776 | 129.947 | 6.047 | | Strong |
| V55 | -110.008 | 6.093 | 127.034 | 6.139 | | Strong |
| A56 | -115.813 | 10.254 | 130.005 | 5.237 | | Strong |
| E57 | -118.648 | 11.456 | 136.519 | 14.876 | | Strong |
| K58 | -121.291 | 16.328 | 140.751 | 15.956 | | Strong |
| T59 | -65.681 | 7.419 | -37.327 | 5.817 | | Strong |
| K60 | -132.440 | 20.024 | 136.233 | 13.056 | | Generous |
| E61 | -124.184 | 15.036 | 135.725 | 12.601 | | Strong |
| Q62 | -97.064 | 16.232 | 122.387 | 6.955 | | Strong |
| V63 | -114.157 | 11.149 | 131.068 | 8.692 | | Strong |
| T64 | -110.600 | 10.200 | 124.483 | 7.768 | | Strong |
| N65 | -112.151 | 10.959 | 127.299 | 6.488 | | Strong |
| V66 | -109.234 | 17.456 | 124.302 | 13.466 | | Strong |
| G67 | 14.051 | 73.475 | -139.744 | 71.042 | | Warn |
| G68 | 81.711 | 7.929 | 3.096 | 11.042 | | Warn |
| A69 | -125.437 | 12.710 | 143.501 | 11.325 | | Strong |
| V70 | -115.294 | 10.058 | 132.898 | 9.069 | | Strong |
| V71 | -107.619 | 11.487 | 129.452 | 9.648 | | Strong |
| T72 | -118.284 | 13.096 | 148.269 | 16.063 | | Strong |
| G73 | -130.477 | 36.326 | 157.726 | 27.206 | | Warn |
| V74 | -106.743 | 12.840 | 129.645 | 6.059 | | Strong |
| T75 | -107.705 | 11.495 | 128.523 | 8.035 | | Strong |
| A76 | -116.511 | 8.989 | 132.137 | 8.385 | | Strong |
| V77 | -116.394 | 11.702 | 131.593 | 7.413 | | Strong |
| A78 | -128.484 | 11.207 | 141.819 | 11.016 | | Strong |
| Q79 | -133.607 | 13.469 | 151.660 | 13.426 | | Strong |
| K80 | -68.025 | 8.461 | -36.877 | 9.913 | | Strong |
| T81 | -135.231 | 13.132 | 146.032 | 15.341 | | Strong |
| V82 | -102.520 | 23.495 | 130.227 | 13.005 | | Strong |
| E83 | -124.728 | 15.787 | 147.703 | 11.645 | | Strong |
| G84 | 121.610 | 48.222 | 25.297 | 107.954 | | Warn |
| A85 | -58.278 | 6.608 | 139.198 | 6.138 | | Strong |
| G86 | 89.999 | 13.008 | -13.013 | 29.224 | | Warn |
| S87 | -77.656 | 20.699 | 150.260 | 10.304 | | Generous |
| I88 | -110.448 | 12.965 | 130.773 | 10.569 | | Strong |
| A89 | -76.151 | 13.834 | 130.183 | 11.812 | | Generous |
| A90 | -121.332 | 12.920 | 132.426 | 9.457 | | Strong |
| A91 | -115.649 | 11.144 | 128.150 | 9.190 | | Strong |
| T92 | -110.073 | 10.685 | 125.243 | 8.893 | | Strong |
| G93 | -145.815 | 41.740 | 169.850 | 33.161 | | Warn |
| F94 | -125.924 | 10.641 | 143.740 | 11.653 | | Strong |
| V95 | -120.102 | 16.073 | 130.717 | 7.520 | | Strong |
| K96 | -115.585 | 9.383 | 130.700 | 8.937 | | Strong |
| K97 | -117.906 | 13.716 | 128.686 | 8.792 | | Strong |
| D98 | -80.464 | 12.082 | 107.998 | 18.186 | | Strong |

^c^Angles are reported in units of degrees (°).

**Table S4:** χ1 angles predicted using TALOS-N

| Residue | Chi1 (χ1) | Δχ1 | | TALOS-N rotamer state | | Rotamer state probability | |
| --- | --- | --- | --- | --- | --- | --- | --- |
| V37 | 175.6 | | 7.5 | | Trans | | 0.771 |
| L38 | -68.4 | | 11.3 | | Gauche - | | 0.865 |
| Y39 | -55.6 | | 8.1 | | Gauche - | | 0.882 |
| V40 | -178.1 | | 10.3 | | Trans | | 0.871 |
| T44 | 58.4 | | 7.8 | | Gauche + | | 0.629 |
| V48 | -175.3 | | 7.9 | | Trans | | 0.637 |
| V49 | -179.3 | | 5.0 | | Trans | | 0.872 |
| V52 | -179.4 | | 7.2 | | Trans | | 0.834 |
| T54 | -60.9 | | 7.8 | | Gauche - | | 0.819 |
| V55 | -177.9 | | 4.3 | | Trans | | 0.872 |
| T59 | -59.4 | | 11.9 | | Gauche - | | 0.671 |
| Q62 | -179.1 | | 10.6 | | Trans | | 0.629 |
| V63 | 178.9 | | 5.1 | | Trans | | 0.872 |
| T64 | -60.8 | | 9.6 | | Gauche - | | 0.819 |
| V66 | -177.1 | | 5.8 | | Trans | | 0.820 |
| V70 | -175.7 | | 6.0 | | Trans | | 0.778 |
| V71 | -173.6 | | 10.6 | | Trans | | 0.872 |
| T72 | 58.7 | | 11.0 | | Gauche + | | 0.738 |
| V74 | 179.5 | | 6.4 | | Trans | | 0.872 |
| T75 | -61.7 | | 10.6 | | Gauche - | | 0.853 |
| V77 | -175.2 | | 5.8 | | Trans | | 0.856 |
| V82 | -178.8 | | 5.4 | | Trans | | 0.856 |
| I88 | -63.2 | | 9.6 | | Gauche - | | 0.693 |
| T92 | -61.0 | | 8.8 | | Gauche - | | 0.819 |
| F94 | -179.3 | | 10.7 | | Trans | | 0.787 |
| V95 | 179.1 | | 7.5 | | Trans | | 0.866 |
| K96 | 179.4 | | 8.8 | | Trans | | 0.745 |

**Table S5:** Unambiguous Manual Distance Restraints used in Xplor-NIH structure calculation^4^.

| **Restraint** | **Distance** | **Spectrum** |
| --- | --- | --- |
| I88CD1-T92CB | 6.00 ± 4.00 Å | 3D ^13^C-^13^C-^13^C with 50 + 500 ms DARR mixing |
| I88CD1-F94CB | 6.00 ± 4.00 Å | 3D ^13^C-^13^C-^13^C with 50 + 500 ms DARR mixing |
| I88CD1-F94CD2 | 6.00 ± 4.00 Å | 3D ^13^C-^13^C-^13^C with 50 + 500 ms DARR mixing |
| I88CD1-F94CZ | 6.00 ± 4.00 Å | 3D ^13^C-^13^C-^13^C with 50 + 500 ms DARR mixing |
| I88CG2-G93C | 6.00 ± 4.00 Å | 3D ^13^C-^13^C-^13^C with 50 + 500 ms DARR mixing |
| I88CG1-K96CA | 6.00 ± 4.00 Å | 3D ^13^C-^13^C-^13^C with 50 + 500 ms DARR mixing |
| I88CG2-F94CG | 6.00 ± 4.00 Å | 3D ^13^C-^13^C-^13^C with 50 + 500 ms DARR mixing |
| I88CB-F94CG | 6.00 ± 4.00 Å | 3D ^13^C-^13^C-^13^C with 50 + 500 ms DARR mixing |
| I88CG1-F94CG | 6.00 ± 4.00 Å | 3D ^13^C-^13^C-^13^C with 50 + 500 ms DARR mixing |
| A90CA-F94CZ | 6.00 ± 4.00 Å | 3D ^13^C-^13^C-^13^C with 50 + 500 ms DARR mixing |
| A91C-F94CZ | 6.00 ± 4.00 Å | 3D ^13^C-^13^C-^13^C with 50 + 500 ms DARR mixing |
| G68C-G93C | 6.00 ± 2.00 Å | 3D ^13^C-^13^C-^13^C with 50 + 500 ms DARR mixing |
| A69CB-G93C | 6.00 ± 2.00 Å | 2D ^13^C-^13^C with 125 ms DARR mixing |
| V74CG2-T92CB | 6.00 ± 4.00 Å | 3D ^13^C-^13^C-^13^C with 50 + 500 ms DARR mixing |
| G68CA-F94CG | 6.00 ± 2.00 Å | 3D ^13^C-^13^C-^13^C with 50 + 500 ms DARR mixing |
| V71CA-G93C | 6.00 ± 2.00 Å | 3D ^13^C-^13^C-^13^C with 50 + 500 ms DARR mixing |
| G47CA-K80CG | 6.00 ± 4.00 Å | 2D ^13^C-^13^C with 500 ms DARR mixing |
| V49CA-A78CB | 6.00 ± 4.00 Å | 2D ^13^C-^13^C with 500 ms DARR mixing |
| G47CA-A78CB | 6.00 ± 4.00 Å | 2D ^13^C-^13^C with 500 ms DARR mixing |
| G47CA-Q79CA | 6.00 ± 4.00 Å | 2D ^13^C-^13^C with 500 ms DARR mixing |
| G47CA-Q79CD | 6.00 ± 4.00 Å | 2D ^13^C-^13^C with 500 ms DARR mixing |
| G47CA-A78C | 6.00 ± 4.00 Å | 2D ^13^C-^13^C with 500 ms DARR mixing |
| G47CA-K80CB | 6.00 ± 4.00 Å | 2D ^13^C-^13^C with 500 ms DARR mixing |
| V49CA-Q79CA | 6.00 ± 4.00 Å | 2D ^13^C-^13^C with 500 ms DARR mixing |
| Y39CZ-T44CA | 6.00 ± 4.00 Å | 2D ^13^C-^13^C with 500 ms DARR mixing |
| K45CE-V48CB | 6.00 ± 4.00 Å | 2D ^13^C-^13^C with 500 ms DARR mixing |
| F94CZ-K96CG | 6.00 ± 4.00 Å | 2D ^13^C-^13^C with 500 ms DARR mixing |
| V37CB-V40CG1 | 6.00 ± 4.00 Å | 2D ^13^C-^13^C with 500 ms DARR mixing |
| E46CB-K80CG | 6.00 ± 2.00 Å | 2D ^13^C-^13^C with 500 ms DARR mixing |
| E46CD-K80CE | 6.00 ± 2.00 Å | 2D ^13^C-^13^C with 125 ms DARR mixing |
| E46CD-K80CG | 6.00 ± 4.00 Å | 3D ^13^C-^13^C-^13^C with 50 + 500 ms DARR mixing |
| E46CG-K80CB | 6.00 ± 4.00 Å | 3D ^13^C-^13^C-^13^C with 50 + 500 ms DARR mixing |

**Table S6:** Summary of simulated annealing protocols in the PASD algorithm.

|  | PASS 2 | PASS 3 | PASS 4 |
| --- | --- | --- | --- |
| Directory on Bessie | 250603_A30P_pass2 | 250608_A30P_pass3 | 250611_A30P_pass4 |
| High Temp I |  |  |  |
| Duration (ps) | 20 | 50 | 50 |
| *k*_lin_ (kcal/mol/Å) | 1 | 0 | 0 |
| *k*_quad_ (kcal/mol/Å^2^) | 0 | 3 | 3 |
| *k*_repul_ (kcal/mol/Å) | 5 | 5 | 5 |
| *Δr_c_* (Å) | ∞ | ∞ | ∞ |
| *w_p_* | 1 | 1 | 1 |
| Number of NOE re-evaluations | 64 | 64 | 64 |
| Number of MC steps | 25 | 25 | 25 |
| *k*_nb_ (kcal/mol/Å^4^) | 0 | 0.004 | 0.004 |
| *s*_nb_ | - | 1.2 | 1.2 |
| Nonbonded interactions | Cα-Cα only | Cα-Cα only | Cα-Cα only |
| *k_R_*_gyr_ (kcal/mol/Å^3^) | 0.001 | 0.001 | 0.001 |
| *k*_dihed_ (kcal/mol/radians^2^) | 150 | 10 | 200 |
| *k*_tDB_ (kcal/mol) | 0.002 | 0.002 | 0.002 |
| High Temp II |  |  |  |
| Duration (ps) | 60 | - | - |
| *k*_lin_ (kcal/mol/Å) | 1 | - | - |
| *k*_quad_ (kcal/mol/Å^2^) | 0 | - | - |
| *k*_repul_ (kcal/mol/Å) | 5 | - | - |
| *Δr_c_* (Å) | 10 | - | - |
| *w_p_* | 0.5 | - | - |
| Number of NOE re-evaluations | 64 | - | - |
| Number of MC steps | 25 | - | - |
| *k*_nb_ (kcal/mol/Å^4^) | 0 | - | - |
| *s*_nb_ | - | - | - |
| Nonbonded interactions | None | - | - |
| *k_R_*_gyr_ (kcal/mol/Å^3^) | 0.001 | - | - |
| *k*_dihed_ (kcal/mol/radians^2^) | 200 | - | - |
| *k*_tDB_ (kcal/mol) | 0.002 | - | - |
| Cooling |  |  |  |
| Duration (ps) | 250 | 250 | 250 |
| *k*_lin_ (kcal/mol/Å) | 1 --> 30 | 0 | 0 |
| *k*_quad_ (kcal/mol/Å^2^) | 0 | 3 --> 30 | 3 --> 30 |
| *k*_repul_ (kcal/mol/Å) | 5 | 5 | 5 |
| *Δr_c_* (Å) | 10 --> 2 | 2.0 --> 0.7 | 2.0 --> 0.7 |
| *w_p_* | 0.5 --> 0 | 0.5 --> 0 | 0.5 --> 0 |
| Number of NOE re-evaluations | 128 | 128 | 128 |
| Number of MC steps | 100 | 100 | 100 |
| *k*_nb_ (kcal/mol/Å^4^) | 0.004 --> 4 | 0.004 --> 4 | 0.004 --> 4 |
| *s*_nb_ | 2.0 | 2.0 | 2.0 |
| Nonbonded interactions | All atoms | All atoms | All atoms |
| *k_R_*_gyr_ (kcal/mol/Å^3^) | 0.001 --> 1.25 | 0.001 --> 1.25 | 0.001 --> 1.25 |
| *k*_dihed_ (kcal/mol/radians^2^) | 200 | 10 --> 300 | 10 --> 300 |
| *k*_tDB_ (kcal/mol) | 0.002 --> 2 | 0.002 --> 2 | 0.002 --> 2 |

**Table S7:** List of Xplor-NIH energy terms used

| Xplor-NIH energy term | Experimental rationale | Restraints per monomer (33-98) | Number of violations in best model | |
| --- | --- | --- | --- | --- |
| NOEPot *(*Interprotomer beta sheet spacing*)* | Define characteristic ~4.7 Å spacing between protomers within a protofilament | - | | 0 |
| NOEPot (unambiguous long-range distance restraints) | Long-range peaks in 2D CC and 3D CCC spectra | 28 | | 0 |
| NOEPot (E46-K80 cross peaks) | E46-K80 salt bridge cross peaks from 2D CC spectrum | 4 | | 1 |
| NOEPot (ambiguous PASD distance restraints) | PASD-assigned distance restraints from 2D CC and 3D CCC spectra | 4187 | | 52 |
| GyrPot | Radius of gyration energy term, keeps overall structure compact | - | | 0 |
| VecPairOrientPot | Vector pair orientation energy term; ensures fibril folds in ribbon-like manner, preventing protomer overlap early on in calculation | - | | 0 |
| RepelPot | Repulsive van der Waals energy term | - | | 84 |
| BOND | Bond lengths | - | | 0 |
| ANGL | Bond angles | - | | 18 |
| IMPR | Bond geometry | - | | 6 |
| CDIH | TALOS-N dihedral angle predictions based on chemical shifts | 127 | | 44 |
| TorsionDBPot | Torsion angle database energy term; bias torsion angles involving heavy atoms towards favorable values observed in high-resolution protein structures | - | | 3 |
| Terminal14Pot | Prevents eclipsed conformations for torsion angles not included in TorsionDBPot | - | | 0 |
